# The Arabidopsis mitochondrial dicarboxylate carrier 2 maintains leaf metabolic homeostasis by uniting malate import and citrate export

**DOI:** 10.1101/2020.04.28.065441

**Authors:** Chun Pong Lee, Marlene Elsässer, Philippe Fuchs, Ricarda Fenske, Markus Schwarzländer, A. Harvey Millar

## Abstract

Malate is the major substrate for respiratory oxidative phosphorylation in illuminated leaves. In the mitochondria malate is converted to citrate either for replenishing tricarboxylic acid (TCA) cycle with carbon, or to be exported as substrate for cytosolic biosynthetic pathways or for storage in the vacuole. In this study, we show that DIC2 functions as a mitochondrial malate/citrate carrier *in vivo* in Arabidopsis. DIC2 knockout (*dic2-1*) results in growth retardation that can only be restored by expressing DIC2 but not its closest homologs DIC1 or DIC3, indicating that their substrate preferences are not identical. Malate uptake by non-energised *dic2-1* mitochondria is reduced but can be restored in fully energised mitochondria by altering fumarate and pyruvate/oxaloacetate transport. A reduced citrate export but an increased citrate accumulation in substrate-fed, energised *dic2-1* mitochondria suggest that DIC2 facilitates the export of citrate from the matrix. Consistent with this, metabolic defects in response to a sudden dark shift or prolonged darkness could be observed in d*ic2-1* leaves, including altered malate, citrate and 2-oxoglutarate utilisation. There was no alteration in TCA cycle metabolite pools and NAD redox state at night; however, isotopic glucose tracing reveals a reduction in citrate labelling in *dic2-1* which resulted in a diversion of flux towards glutamine, as well as the removal of excess malate via asparagine and threonine synthesis. Overall, these observations indicate that DIC2 is responsible *in vivo* for mitochondrial malate import and citrate export which coordinate carbon metabolism between the mitochondrial matrix and the other cell compartments.

**SIGNIFICANCE STATEMENT:** Mitochondria are pivotal for plant metabolism. One of their central functions is to provide carbon intermediates for the synthesis of critical building blocks, such as amino acids. Malate import and citrate export are two of the most recognised and specialised features of the mitochondrial role in the plant cellular metabolic network, yet the possibility that a single carrier would unite both functions has not been considered. Here, we have demonstrated that DIC2 preferentially fulfils these two functions in *Arabidopsis thaliana in vivo*, making it a bifunctional gateway for two major metabolite fluxes into and out of the mitochondrial matrix in the plant cell. Our results highlight the significance of DIC2 in cooperation with other mitochondrial carriers in maintaining metabolic balance even under challenging environmental conditions.

## INTRODUCTION

Malate is a prominent metabolite that occupies a pivotal node in the regulation of plant carbon metabolism. It is the mainstay of leaf respiration and metabolic redox shuttling between organelles (1). Early studies with isolated plant mitochondria demonstrated that exogenous malate can be translocated by an unknown mechanism into the mitochondrial matrix where it is then rapidly oxidized by both mitochondrial malate dehydrogenase (mMDH) and malic enzyme (NAD-ME), generating oxaloacetate (OAA) and pyruvate as products (2, 3). OAA inhibits mitochondrial respiration by direct inhibition of succinate dehydrogenase and due to the fact that the chemical equilibrium of the mMDH reaction highly favours OAA conversion to malate, thereby diverting NADH away from the electron transport chain (ETC) (4, 5). The NAD-ME activity prevents OAA accumulation in mitochondria, and thus allows the continued oxidation of malate produced in concert with phosphoenolpyruvate carboxylase and malate dehydrogenase activities in the cytosol (6–8), even in the absence of glycolysis-derived pyruvate. Both experimental evidence and flux-balance model predictions indicate that TCA cycle-driven respiration is largely inhibited in the light and that mMDH generally operates in the reverse direction to facilitate redox coupling, resulting in net mitochondrial OAA import and malate efflux (9–13). At night, the requirement for exchanging reducing equivalents between mitochondria and chloroplasts via a malate valve (1, 14, 15) is believed to be minimal due the cessation of both photorespiration and photoinhibitory conditions. The mitochondrial NAD-ME activity and its transcript abundance is highest in the dark (16), therefore the synthesis and oxidation of malate is expected to be carried out by both mMDH and NAD-ME to support the synthesis of ATP (17). These metabolic conditions would enable maximal citrate oxidation in the mitochondrial matrix (18, 19), with excess citrate being exported for storage in the vacuole (20). Inhibition in the synthesis of citrate from malate-derived OAA in mitochondria and its export leads to floral sterility in potato plants (21) and a change in nitrogen incorporation into amino acids and metabolism in tomato leaves (22). Defects in controlling mitochondrial malate use cause different phenotypic changes and metabolic remodelling: the absence of mMDH activity in Arabidopsis results in a slow growth phenotype and an elevated leaf respiration rate (23), and the loss of NAD-ME in Arabidopsis causes a significant diversion of excess malate to amino acid synthesis at night (16). Even though the mitochondrial transport of malate and citrate is clearly at the heart of the remarkable metabolic flexibility of plants, the identity of the transporter(s) that underpin those fluxes *in vivo* has remained inconclusive.

By identifying homologues of yeast and mammalian carriers in *Arabidopsis thaliana*, several possible candidates that may contribute to mitochondrial malate transport in plants were identified (24–26). In contrast to the historical, well-established models that mitochondrial metabolite transporters are discrete and highly specific (27–29), these carriers appeared to lack substrate specificities under *in vitro* conditions – for example, dicarboxylate carrier (DIC) isoforms in proteoliposomes can rapidly exchange sulfate with phosphate, malate, OAA and succinate, with an apparent low exchange activity in the presence of citrate, 2-oxoglutarate and fumarate (24). Although *in vitro* studies have been instrumental to reveal what transport activities can be mediated by a protein, they have limitations that turn out to be critical in the case of plant mitochondrial organic acid transporters. For instance, they cannot suitably consider the – often unknown and probably highly changeable – relative metabolite pool sizes and fluxes that the transporters face *in vivo*. Reconstitution with a specific orientation of the transporter is not possible in most *in vitro* systems, but critical for substrate specificity and respiratory physiology *in vivo*. Further, the specific local lipid environment and the pronounced electrochemical gradients (Δψ, ΔpH), both of which are likely to be central for inner mitochondrial membrane transport activity and specificity (17), can only be roughly considered. As such, additional information is needed on the function of any transporter candidate within a physiologically more meaningful context. Yet, *in planta* studies of mitochondrial metabolite transporter function that address the question of their *in vivo* function, requiring a systems perspective to consider their integration in the metabolic network, have been lacking.

In this study, we reveal a primary role for DIC2 in malate import and citrate export from Arabidopsis mitochondria. Through a reverse genetic approach in combination with comprehensive *in vitro*, *in organello*, and *in vivo* analyses, we obtained strong evidence that plant mitochondrial malate and citrate exchange as mediated by DIC2 plays a critical role in dynamically coordinating anaplerotic metabolism with assimilatory and catabolic pathways between the mitochondria and other cellular compartments.

## RESULTS

### DIC2 mutation causes decelerated vegetative growth rate that cannot be compensated by other mitochondrial dicarboxylate carrier isoforms

DIC2 is targeted to the mitochondrion when transiently expressed (30) and has recently been identified in the mitochondrial proteome by mass spectrometry (31, 32). When DIC2-GFP is stably expressed in Arabidopsis, the fusion protein was found as spherical to elongated structures that colocalised with the mitochondrial dye tetramethylrhodamine (TMRM) (Fig. S1). No overlapping of signal between DIC2-GFP and chloroplast auto-fluorescence was observed in leaf tissue, indicating that DIC2 localises to mitochondria in Arabidopsis.

Three T-DNA insertion lines were obtained in order to study the possible physiological and metabolic consequences of a defect in DIC2 (At4g24570) function *in planta*. Only the homozygous *dic2-1* line has a T-DNA insertion within the open reading frame of *DIC2*, resulting in the complete loss of *DIC2* transcript (Fig. 1A and Fig. S2A). Using selective reaction monitoring mass spectrometry (SRM-MS) to determine protein abundances (33), we found that two different DIC2 peptides, detectable in Col-0, were not detectable above baseline chemical noise in mitochondria isolated from *dic2-1* plants (Fig. 1A and Fig. S2B), confirming *dic2-1* as a null mutant for DIC2. Given that *DIC2* is more expressed in green tissues than in roots (24), it is expected that the loss of DIC2 would have a greater impact on vegetative growth. Indeed, the homozygous *dic2-1* was characterized by a significantly decreased rate of rosette expansion (Fig. 1B and Fig. S2C). From day 42, the number of leaves emerged from Col-0 and *dic2-1* was identical, but the mutant leaves were smaller and curly with a more rugose surface (Fig. S2D). *dic2-1* could not reach the full rosette diameter of Col-0 after one week of bolting (Fig. S2E). *dic2-1*-specific phenotypes could be observed regardless of the photoperiod (Fig. S2F). These phenotypes could be restored by introducing *DIC2* under the control of its native promoter (*dic2-1/gDIC2*) or a cauliflower mosaic virus 35S promoter (*dic2-1/DIC2 OE*). While full complementation was reached through the native promoter, 35S-driven expression of *DIC2* resulted in a slight delay in phenotype restoration in the *dic2-1* background and led to a reduced rosette size in Col-0 background (Fig. 1B, Fig. S2G and Table S1), indicating that the correct dose of *DIC2* expression is essential for normal leaf growth.

**Figure 1.**
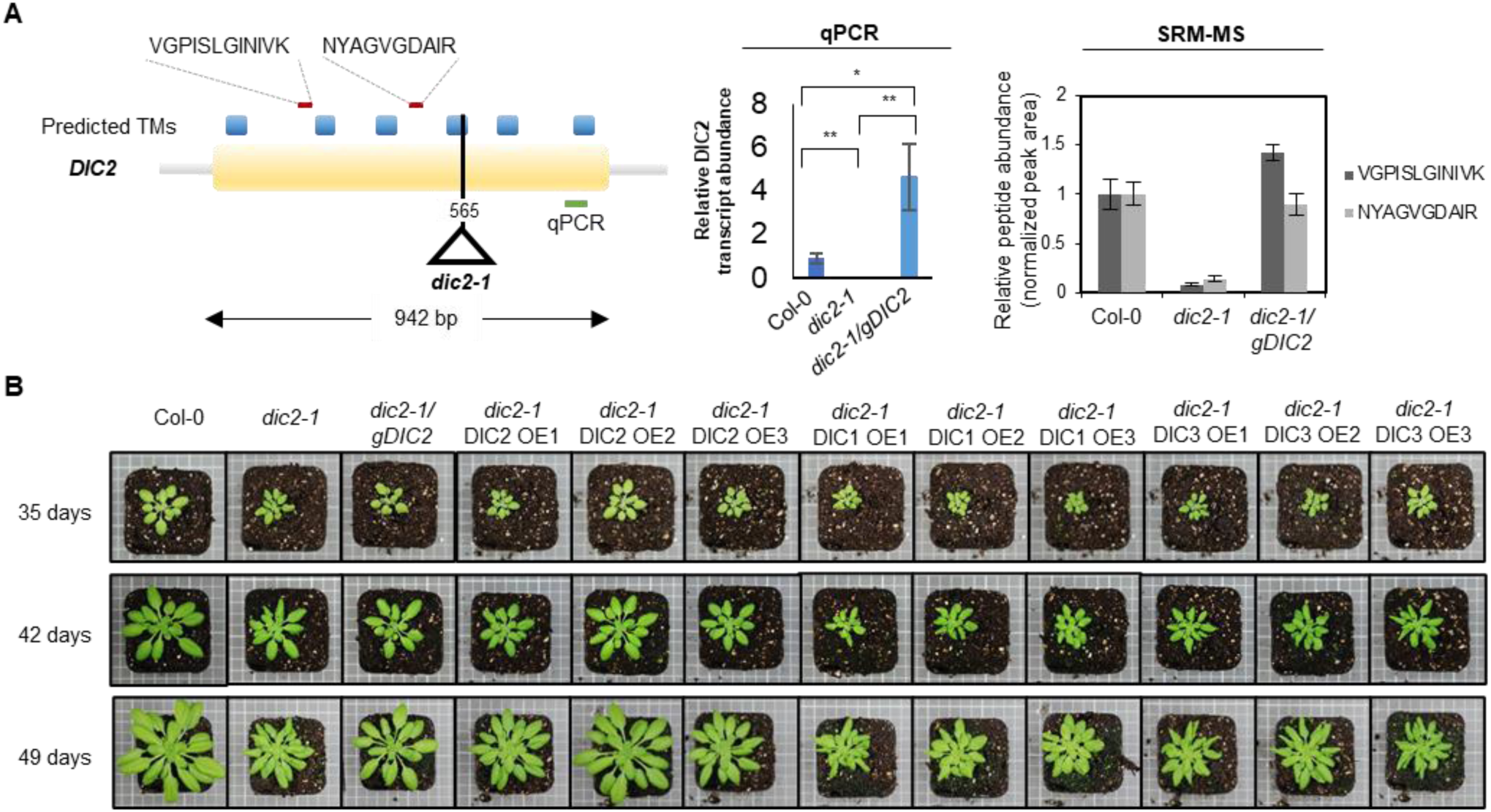
Phenotypic characterization of the *DIC2* mutant. (A) Left, the gene model of DIC2 showing predicted transmembrane domains based on ARAMEMNON consensus prediction (95), the position of T-DNA insertion in the *dic2-1* line and locations of peptides for LC-MRM-MS (in red lines) and the transcript for qPCR (in green line). Middle, expression levels of *DIC2* as determined by qPCR in different genotypes (n = 4). Right, LC-MRM-MS abundance analysis of unique peptides of DIC2 using the quantifier ion transitions VGPISLGINIVK and NYAGVGDAIR (n = 3). Mean ± S.E., with asterisks denote significant differences between *mcc-1* vs Col-0 and *mcc-1* vs *mcc-1/gMCC* based on ANOVA and Tukey’s post-hoc analysis (* p < 0.05; ** p < 0.01). (B) Vegetative phenotype of Col-0, *dic2-1* and complemented lines (*gDIC2*, native promoter; *OE*, 35S promoter) grown on soil under short day condition (8 h light/16 h dark) on 35, 42 and 49 days after germination. Representative top view of various genotypes is shown. Expression levels are shown in Table S1.

DIC2 shares high amino acid sequence similarity as well as broad and identical substrate preferences with two mitochondrial dicarboxylate carriers DIC1 and DIC3 as determined *in vitro* (24). Liposome-based transport studies also found overlapping substrate preferences with succinate-fumarate carrier 1 (SFC1), dicarboxylate-tricarboxylate carrier (DTC) and uncoupling protein 1 (UCP1) (25, 26, 34). The loss of DIC2 resulted in a modest two-to three-fold increase in *DIC1* and *DIC3* transcripts at night, but not at mid-day, while the transcripts of the other three proteins were not affected (Fig. S2H). To test the hypothesis of functional redundancy within the DIC family, we attempted to rescue the *dic2-1* phenotype by expressing *DIC1* and *DIC3* in the *dic2-1* background (*dic2-1*/*DIC1 OE* and *dic2-1*/*DIC3 OE* respectively), but neither could complement the loss of DIC2, not even partially (Fig. 1B). These results indicate that DIC2 is unlikely to share substrate preferences and/or specificities with DIC1 or DIC3 *in planta* and emphasize the need for an *in vivo* assessment.

### Malate is the major dicarboxylate substrate of DIC2 in isolated Arabidopsis mitochondria

The inability of DIC1 or DIC3 to rescue *dic2-1* phenotypes prompted us to reinvestigate DIC2 function in detail. In order to maintain DIC2 close to its true functional context, while being able to control substrate availabilities, we used isolated mitochondria as a model system to connect previous *in vitro* insights from liposome assays and the more complex *in vivo* situation. We first examined if malate and succinate are the main substrates for DIC2 in isolated mitochondria, as inferred previously by a liposome-based approach (24). To this end, mitochondria purified from Col-0, *dic2-1* and *dic2-1/gDIC2* were tested for their ability to consume dicarboxylates under state III respiration conditions (measured as O_2_ consumption coupled to ATP synthesis) and to generate and export metabolites synthesised from the substrates provided (measured by LC-SRM-MS analysis of the separated mitochondrial and the extra-mitochondrial space fractions; Fig. 2A).

**Figure 2.**
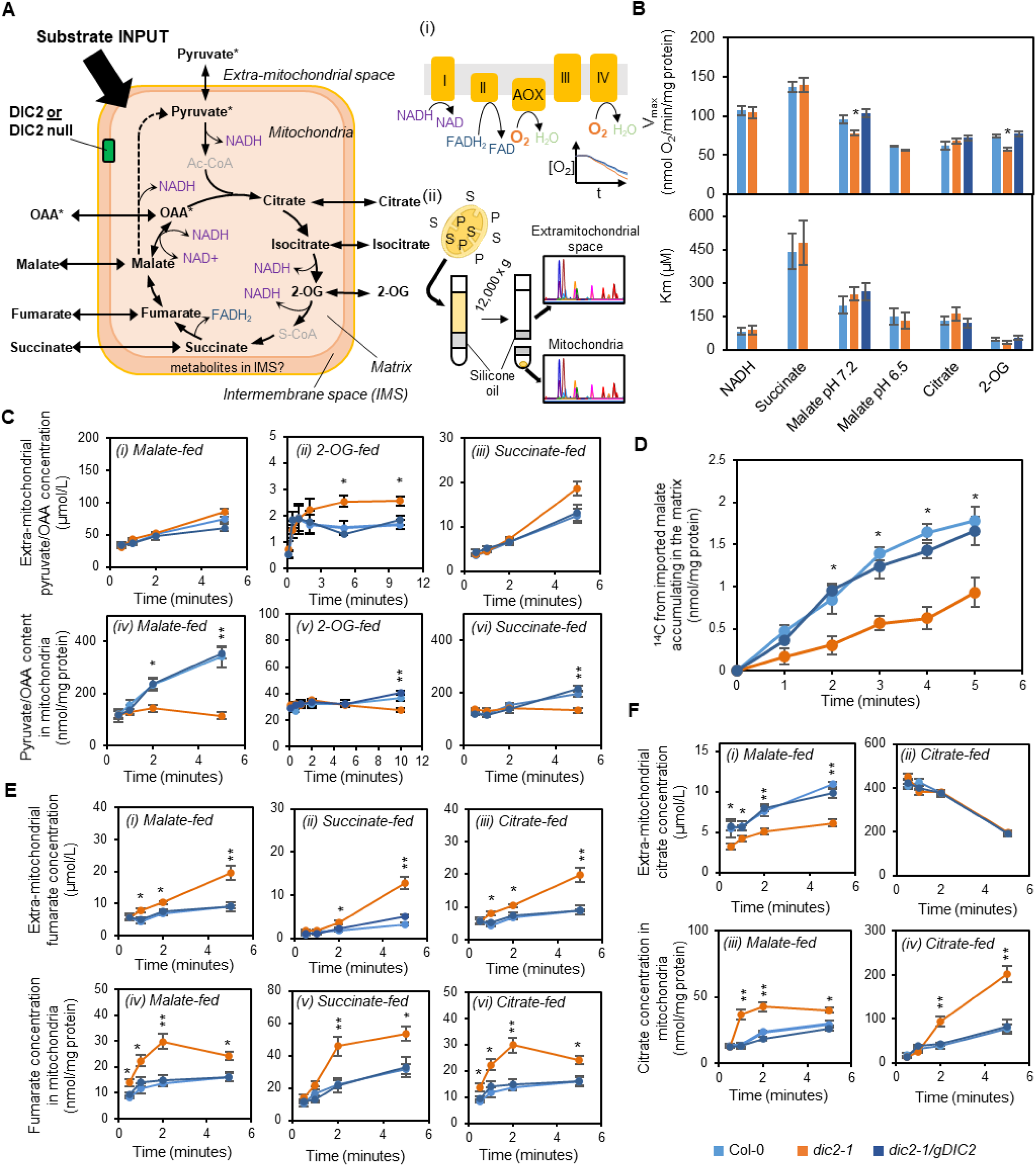
Uptake, consumption and export of TCA cycle intermediates by isolated mitochondria. (A) Experimental design for monitoring substrate consumption, product formation and metabolite transport kinetics of isolated mitochondria. On the left, the mitochondrial TCA cycle and pyruvate metabolising and generating steps are shown, along with the steps that generate reductants for consumption by the ETC and the movements of organic acids across the membranes. Note that the feeding substrate, which is in excess, could cause inhibition of specific steps of the TCA cycle (e.g. feedback inhibition of citrate synthase by citrate). (i) Measurement of oxygen consumption by substrate-fed mitochondria using Clarke-type oxygen electrode is an indirect assay for simultaneously measuring substrate uptake and consumption and subsequent transfer of reductant to the ETC. (ii) In a second approach, mitochondria are fed with a substrate under energised condition. After indicated time interval, mitochondria are separated from extra-mitochondrial space by centrifugation through a silicone oil layer. These fractions are collected, and substrates (S) and products (P) are quantified by SRM-MS. (B) Bar graphs of V_max_ (upper panel) K_m_ (lower panel) for O_2_ consumption in the presence of substrate indicated. Mean ± S.E. (n ≥ 5). (C) Time-courses of pyruvate and/or OAA concentrations in extra-mitochondrial space (i-iii) and the matrix (iv-vi) incubated with the indicated feeding substrate. Mean ± S.E. (n = 4). (D) [^14^C]-malate uptake into isolated mitochondria as a function of time under non-energising conditions. Isolated mitochondria were incubated in minimal transport buffer containing 200 µM malate without energising cofactors for the time indicated. Mean ± S.E. (n = 4). (E) Time-courses of fumarate concentrations in the extra-mitochondrial space (i-iii) and matrix (iv-vi) incubated with the indicated feeding substrate. Mean ± S.E. (n = 4). (G) Time-courses of citrate concentrations in the extra-mitochondrial space (i-ii) and matrix (iii-iv) incubated with the indicated feeding substrate. Mean ± S.E. (n = 4). Asterisks denote significant differences between *mcc-1* vs Col-0 and *mcc-1* vs *mcc-1/gMCC* based on ANOVA and Tukey’s post-hoc analysis (* p < 0.05; ** p < 0.01). Response curves shown in C, E and F can also be found in Fig. S5.

Titrations with respiratory substrates at different concentrations revealed no obvious differences in the rate of internal or external NADH-driven O_2_ consumption between Col-0 and *dic2-1* (Fig. 2B; Fig. S3A). No change in the abundance or activity of electron transport chain (ETC) supercomplexes (Fig. S4A), the relative abundance of TCA cycle enzymes, pyruvate dehydrogenase or NAD-ME (Fig. S4B), or their K_m_ and V_max_ (Table S2) were observed when mitochondria isolated from these genotypes were compared. This eliminated the possibility of a clear defect in the ETC or a specific step of the TCA cycle in *dic2-1*; instead, any shift in the distribution of specific metabolites between the matrix and extra-mitochondrial space in the mutant could reasonably be attributed directly to the absence of DIC2 and/or indirectly to the enzymatic regulatory compensation as the result of DIC2 loss.

Through the action of the TCA cycle and NAD-ME, imported substrates can generate both OAA and pyruvate (Fig. 2A). Note that it is difficult to experimentally assess OAA and pyruvate independently because OAA undergoes spontaneous decarboxylation into pyruvate; therefore, we considered the pyruvate signal to represent both pyruvate and OAA levels. Upon feeding externally with 500 µM malate, pyruvate and/or OAA abundance increased linearly over time in mitochondria of Col-0 and *dic2-1/gDIC2*, whereas it levelled off in *dic2-1* (Fig. 2C, panel iv). The magnitude of *dic2-1*-specific increase in pyruvate and/or OAA content in mitochondria was far less drastic for succinate or 2-OG feeding, both of which led to the formation of malate within the matrix (Fig. 2C, panel v-vi). Consistent with this, *dic2-1* mitochondria exhibited a 20–30% lower V_max_ for malate-dependent state III O_2_ consumption at pH 7.2, although the K_m_ was unchanged (Fig. 2B). When malate-dependent O_2_ consumption by isolated mitochondria was measured at the optimal pH for NAD-ME activity (pH~6.5)(35), no difference in K_m_ and V_max_ between *dic2-1* and Col-0 was observed. These data pinpoint reduced malate availability in the matrix of malate-fed *dic2-1* mitochondria as a plausible cause for the observed decrease in malate-dependent respiration (via mMDH) and pyruvate and/or OAA formation. To test more directly whether DIC2 indeed imports malate into mitochondria and whether this import is restricted in *dic2-1*, we monitored the transport activity for ^14^C-malate by isolated mitochondria in a reaction medium that lacked cofactors and ADP to allow malate to accumulate while preventing its conversion into other TCA cycle intermediates. Under such conditions, the initial uptake rate of 200 µM [^14^C]-malate into mitochondria of *dic2-1* was three-fold lower compared to Col-0 and *dic2-1/gDIC2* (Fig. 2D). Hence, DIC2 possesses a significant malate uptake capacity in isolated Arabidopsis mitochondria even in the presence of other potential carriers for malate transport.

If the role of DIC2 in 2-OG and/or succinate uptake is as important as malate transport, mutant mitochondria would be expected to display a lower O_2_ consumption rate in their presence as well as to accumulate and/or export less of their nearest TCA cycle product(s) when these substrates are supplied. There was no significant change in succinate-stimulated Complex II-linked O_2_ consumption (Fig. 2B), so the mutant growth phenotype was unlikely to be caused by a reduced mitochondrial succinate uptake and oxidation. 2-OG-dependent state III respiration in *dic2-1* mitochondria was reduced in *dic2-1* mitochondria, to an extent which was strikingly similar to malate oxidation at pH 7.2 (Fig. 2B). However, DIC2 is unlikely to be a major 2-OG importer, since 2-OG, malate and succinate levels in *dic2-1* mitochondria and extramitochondrial space were not altered when 2-OG was supplied (Fig. S5C). This anomaly between O_2_ consumption and substrate accumulation patterns may be explained by a partial feedback inhibition of mMDH reducing NADH necessary for the ETC activities and resulting in a flux diversion of malate to fumarate (Fig. 2A). Consistent with this we observed increases in fumarate accumulation and efflux by *dic2-1* mitochondria when supplied with malate and succinate (Fig. 2E, panel i-ii and iv-v). In addition, it could be due to the observed increase in pyruvate and/or OAA export rate to relief the product inhibition of mMDH when supplied with 2-OG (Fig. 2C, panel ii).

### DIC2 is one of the mitochondrial carriers that determine the fate of citrate in energised mitochondria

The other main discovery from our substrate feeding experiments was a much higher amount of citrate accumulating inside *dic2-1* than Col-0 mitochondria after malate (Fig. 2F panel iii), succinate or 2-OG feedings (Fig. S5C-D). If OAA and/or pyruvate exports were also limited in *dic2-1* mitochondria, matrix pyruvate and/or OAA contents would be expected to accumulate more over time with a slower combined rate of their export to the extramitochondrial space. However, pyruvate and/or OAA appear to be maintained in the mutant mitochondria in the same way as in Col-0 after malate or succinate feedings (Fig. 2C, panel i and iii). In contrast, the rate of export of citrate to the extra-mitochondrial space was decreased in *dic2-1* mitochondria when compared to Col-0 and *dic2-1/gDIC2* in malate (Fig. 2F panel i), succinate or 2-OG feedings (Fig. S5C-D). The degree of decrease in each case was different depending on substrate concentration and the number of subsequent enzymatic steps leading to citrate formation. We hypothesised that excess citrate in *dic2-1* mitochondria was caused by a defect in citrate export which then prevented OAA condensation in the citrate synthase reaction and thus suppressed mMDH activity (36). To confirm this, we tested the ability of mitochondria to directly consume citrate (Fig. S5E). Under such conditions, NAD-ME and mMDH activities became inhibited, as demonstrated by the low mitochondrial export rate of pyruvate and/or OAA (Fig. S5E). There was no change in citrate-dependent O_2_ consumption mitochondria from all genotypes (Fig. 2B), further confirming that citrate import was not affected and the reduction in malate- and 2-OG-dependent respiration by *dic2-1* was likely caused by the citrate accumulation in the matrix triggering an OAA inhibition of mMDH.

While the uptake rates of citrate by mitochondria were not altered significantly over time (Fig. 2F, panel ii), the rate of citrate accumulation in *dic2-1* mitochondria was two- to three-fold higher than Col-0 or *dic2-1/gDIC2* (Fig. 2F, panel iv). *dic2-1* mitochondria appeared to be oxidising excess citrate through the TCA cycle (Fig. 2A) as evident by a higher rate of 2-OG and fumarate accumulation in *dic2-1* mitochondria and an increased fumarate export (Fig. 2E, panel iii and vi; Fig. S5E). Similar results were also observed when mitochondria were externally provided with isocitrate (Fig. S5F). We could not observe any obvious change in import and export rates of succinate, 2-OG and pyruvate by *dic2-1* mitochondria, indicating that other carriers with a similar set of *in vitro* substrates as DIC2, as previously observed in proteoliposome-based studies, may be able to compensate for its absence. Taken together, DIC2 is directly responsible for malate uptake, and we hypothesised that either citrate is the counter ion, or DIC2 is able to separately transport these organic acids in opposing directions. The fact that citrate export was not completely abolished by the loss of DIC2 would suggest that there are also other citrate export carriers present in the inner mitochondrial membrane.

### DIC2 function effects leaf dark respiration by modulating NAD homeostasis

We then set out to determine if defects in mitochondrial malate import and citrate export operate *in planta* and if they could explain the observed phenotypes of *dic2-1*. In total we investigated three scenarios where leaf mitochondria operate under different flux modes: in the light, during the transition from light to dark and in the dark. We first carried out chlorophyll fluorescence and infrared gas-exchange analyses at different light intensities, because changes in photosynthetic performance can cause many metabolic perturbations and mask other primary metabolic effects in mutants. There was no change in the CO_2_ assimilation rate (Fig. 3A), stomatal conductance, transpiration rate, photosynthetic electron transport rate, photosystem II quantum yield or nonphotochemical quenching in *dic2-1* leaves (Fig. S6A-E). When plants were shifted to high light for 3- and 16-h, we did not detect any changes in the maximum photochemical efficiency of PSII (F_v_/F_m_) between *dic2-1*, Col-0 and *dic2-1/gDIC2* (Fig. S6F). These results indicated that the loss of DIC2 has no direct impact on photosynthetic performance.

**Figure 3.**
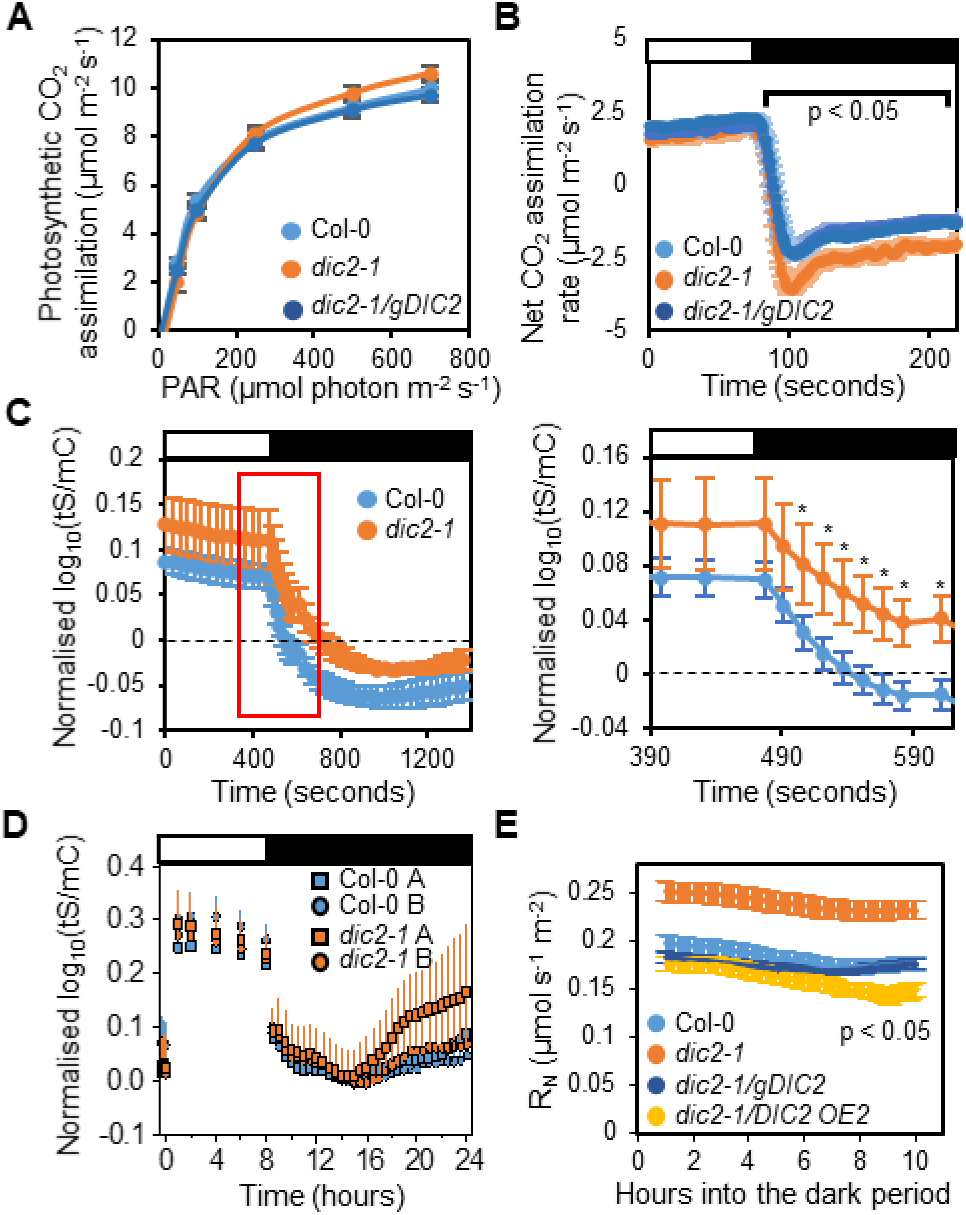
Photosynthetic and respiratory phenotypes of DIC2 knockout. (A) Photosynthetic CO_2_ assimilation rate at different photosynthetic active radiation (PAR) with CO_2_ concentration at 400 p.p.m. and temperature at 22°C (n = 6, mean ± S.E., no significant difference based on one-way ANOVA). (B) Determination of post-illumination CO_2_ burst. A single leaf illuminated with actinic light of 1000 μmol m^−2^ s^−1^, CO_2_ concentration of 100 p.p.m. at 25°C was darkened for two minutes and post-illumination burst was monitored in the first 30 s (n = 6, mean ± S.E., data points within the bracket indicate significant differences of p < 0.05 based on one-way ANOVA Tukey post-hoc test). (C) Cytosolic NAD redox dynamics of 6-week-old leaves in response to a sudden light-to-dark transition. Redox changes of the NAD pool correlate to the Peredox-mCherry ratio (log_10_(tS/mC)), i.e. high ratio indicates high NADH/NAD^+^ ratio. Dark adapted leaf discs were illuminated at actinic light of 220 μmol m−2 s−1 according to *SI Methods* before they were transferred to the dark. The 5-min light/15-min dark time course is shown on the left, with the red rectangle indicating the moment when significant differences in NAD redox state were observed. Zoom-in of this red rectangle time interval is shown on the right. Each data point represents mean ± S.D. (n ≥ 6), with asterisks indicate a significant difference with p < 0.001 based on multiple t-tests (alpha = 5%) from the Holm-Sidak Method. (D) Changes in the cytosolic NAD redox state over 8-h light/16-h dark diurnal cycle. Leaf discs were exposed to actinic light of 120 μmol m−2 s−1 before light was switched off. Two independent lines (A and B) for each genotype were measured. Data shown indicate mean ± SD (n ≥ 8) with no significant differences found between lines. (E) Time course of leaf respiration measurements in the dark (RN) as measured by Q2 (n > 8, mean ± S.E., all time points are significantly difference with p < 0.05 based on one-way ANOVA Tukey post-hoc test). Response curves shown in C, E and F can also be found in Fig. S5.

We next examined the transition from light to darkness where leaf mitochondria undergo several rapid changes in metabolic activity as photorespiration ceases and non-cyclic mode of the mitochondrial TCA cycle is activated. Upon transfer of illuminated leaves to darkness, *dic2-1* exhibited a more rapid release of CO_2_ in the first 30-40 s before reaching a higher rate of steady-state respiratory CO_2_ release compared to Col-0 and *dic2-1/gDIC2* (Fig. 3B). The sharp CO_2_ release upon sudden darkness is indicative of a post-illumination burst (PIB) of respiration, an estimate of the degree of respiratory glycine oxidation (37). To examine if photorespiratory activities were altered in the mutant, we grew the mutant under high CO_2_ (0.2%) to suppress photorespiration but no rescue of the phenotype was achieved (Fig. S6G). These data indicate that an altered photorespiratory activity after dark shift is not the primary cause for the change in PIB response by *dic2-1*. Since malate/OAA utilisation in mitochondria can control the degree of glycine oxidation by regulating NADH/NAD^+^ levels (e.g. via a malate shuttle) (4), we hypothesised that a PIB increase in *dic2-1* could be linked to altered distribution of TCA cycle metabolites between mitochondria and cytosol (and possibly plastids), leading to changes in metabolic and/or NAD redox state in these compartments. To capture any rapid and transient changes in NAD redox state during a sudden dark shift, we utilised a fluorescent protein biosensor Peredox‐mCherry which reports cytosolic NADH/NAD^+^ through the ratio of tSapphire to mCherry fluorescence (log_10_(tS/mC)), where a higher log_10_(tS/mC) corresponds to a more reduced NAD pool (38, 39). We observed equally high NADH/NAD^+^ ratios in Col-0 and *dic2-1* plants during illumination (Fig. 3C). However, upon transfer to darkness, the expected decline in NADH/NAD^+^ ratios in the first 100 seconds was significantly slower in *dic2-1* compared to Col-0. Both differences in PIB and NAD redox status upon rapid transition to darkness indicate some defect in switching between light and dark flux modes by *dic2-1* mitochondria.

Thirdly, we examined metabolic changes to *dic2-1* leaves during the night. DIC2 expression steadily increased over the course of night-time, followed by a decline from the peak at the end of dark photoperiod to the lowest level in the light (Fig. S6H). Intriguingly, while there was no difference in the NADH/NAD^+^ ratio between the two genotypes over the course of the dark-light-cycle (Fig. 3D), the leaf night time respiration rate (RN) remained consistently higher in the mutant (Fig. 3E). Given that primary metabolites are crucial determinants for regulating transcript abundance during a diurnal cycle (40), there appears to be a metabolic rearrangement in the mutant to avoid any detrimental impact of DIC2 absence on NAD redox equilibrium, albeit at the expense of heightened respiration (Fig. 3B and 3E) and slower vegetative growth (Fig. 1B). Taken together, DIC2 activity may have a greater influence on metabolism and mitochondrial respiration in the dark, most likely through modulating the TCA cycle steps in the cytosol and/or mitochondria to maintain metabolic homeostasis required for fundamental cellular functions.

### 2-oxoglutarate sharply accumulates in *dic2-1* leaves upon shift to darkness

To gain further insight into how DIC2 transport functions are integrated into metabolism and influences NAD redox state *in vivo*, we carried out metabolite profiling of 6-week-old leaf samples collected at different time points of a diurnal cycle (Fig. 4A and Fig. S7). The metabolite profiles of malate, fumarate and succinate over a diurnal cycle were generally similar between genotypes. Also, a similar pattern in daytime glycine and serine accumulation was observed, despite a clear difference in PIB (Fig. 3B), further confirming that DIC2 function does not directly affect photorespiratory flux. The most notable metabolite level changes in *dic2-1* plants compared to Col-0 and *dic2-1/gDIC2* were observed in the first-hour after the sudden shift to darkness, including two- to four-fold higher abundances of pyruvate, citrate and isocitrate, and a remarkable 10-fold increase in 2-oxoglutarate abundance (2-OG). Several amino acids linked to 2-OG utilization also changed in abundance in *dic2-1*. In particular, the glutamate pool in *dic2-1* was larger than Col-0 and *dic2-1/gDIC2* plants one hour after the dark shift and during the light photoperiod, while GABA and glutamine abundances remained unchanged. Aspartate and alanine, which can be converted into OAA and pyruvate respectively in a reversible reaction that requires glutamate/2-OG transamination, were also more abundant in *dic2-1* after one hour of darkness, while aspartate-derived threonine accumulated in the mutant throughout the diurnal cycle. Branched chain amino acids (BCAAs), which use 2-OG as co-substrate to initiate their catabolism in mitochondria (41), were generally higher in abundances in *dic2-1* plants at night. Overall, the metabolite profiles revealed that the loss of DIC2 function causes an impaired 2-OG metabolism in the dark, which may provide a metabolite buffer to compensate for a defect in mitochondrial DIC2 transport activity while cytosolic NAD redox state is maintained.

**Figure 4.**
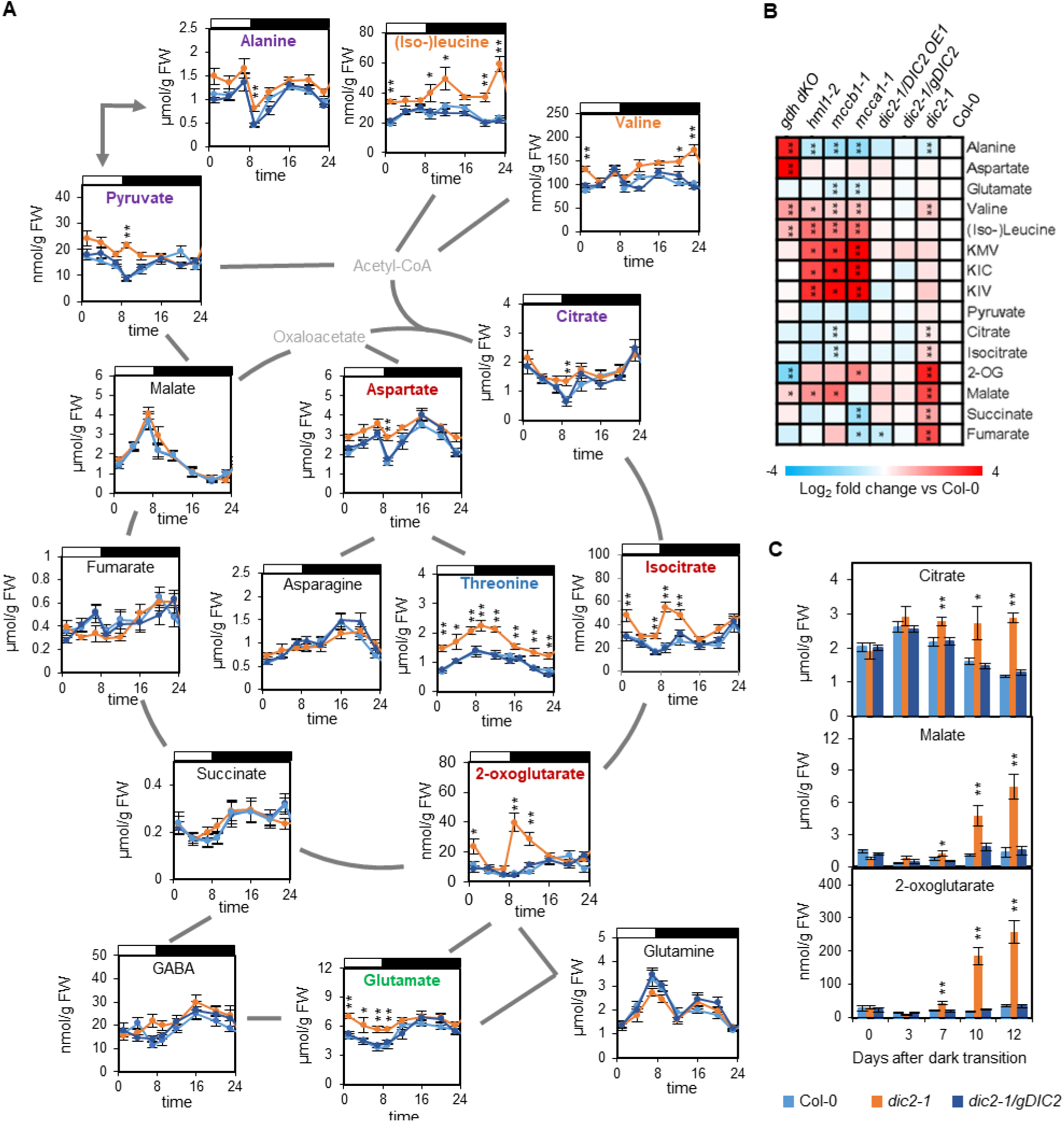
Quantitative analysis of metabolites associated with the TCA cycle in a diurnal cycle and during prolonged darkness. (A) Plants were grown under short day conditions for 6 weeks and leaf discs were collected at 1, 4, 8, 12 and 15 hours after dark shift and 1, 4 and 7 hours after light shift. Metabolites in this figure were analysed by LC-MRM-MS. Metabolites are coloured according to their accumulation pattern in *dic2-1* in a diurnal cycle: Orange, accumulates at night; Red, accumulates during light and dark shift; Green, accumulates predominantly during the day; Blue, accumulates throughout a diurnal cycle; Purple, accumulates only after the first hour of dark shift; Grey, metabolite not measured. Each data point represents mean ± S.E. (n ≥ 6), asterisks indicate a significant change as determined by the one-way ANOVA with Tukey post-hoc test (* p < 0.05; ** p < 0.01). (B) Heat map showing log2-fold change, relative to Col-0, of TCA cycle intermediates, selected amino acids and branched chain amino acid derivatives in four-week-old transgenic lines treated with 10 days of prolonged darkness (n = 7). BCKAs inclue: KIV, 2-oxoisovalerate; KIC, 2-oxoisocaproate; KMV, 2-oxo-3-methylvalerate. * p < 0.05 and ** p < 0.01 according to the Student t-test. (C) Changes in citrate, 2-OG and malate content in Arabidopsis plants after 0, 3, 7, 10 and 12 days of extended darkness treatment (n = 7). Each data point represents mean ± S.E. Asterisks indicate a significant change as determined by one-way ANOVA with Tukey post-hoc test (* p < 0.05; ** p < 0.01).

### A defect in organic acid use causes accelerated leaf senescence of the DIC2 knockout during prolonged darkness

Elevated level of BCAAs in *dic2-1* (Fig. 4A) may indicate a possible defect in mitochondrial BCAA catabolism due to changes in the uptake of BCAAs and/or their derivatives into the matrix. Most BCAA catabolism mutants accumulate BCAAs and are more susceptible to early senescence during prolonged darkness (42–47). To test if intermediates of BCAA catabolism could also be DIC2 substrates, we subjected Arabidopsis plants to darkness for 15 days. The mutant exhibited an accelerated decline in F_v_/F_m_ beginning around 7 days of darkness (Fig. S8A). Leaves of *dic2-1* were more yellowed compared with the Col-0 and the *dic2-1/gDIC2* after 12 days (Fig. S8B). When 15 days dark-treated plants were transferred back to the normal short-day cycle, only Col-0 and *dic2-1/gDIC2* could recover after seven days. Next, we measured the abundance of organic acids, amino acids and branched chain 2-oxoacids (BCKA) in 10 days dark-treated transgenic plants with known mitochondrial responses to carbon starvation (Fig. 4B). Knockout mutants with defects in mitochondrial BCAA catabolism, *mcca1-1, mccb1-1* and *hml1-2*, displayed accelerated senescence and had higher amounts of BCAAs than Col-0 as observed previously (43, 48), as well as accumulated BCKA. In comparison, the double knockout of glutamate dehydrogenase (*gdh dKO*), which does not directly participate in BCAA catabolism but senesces more rapidly under prolonged darkness (49), significantly accumulated BCAAs without altering the abundance of BCKA. The loss of DIC2 resulted in slight but significant accumulation of valine but not leucine, isoleucine and BCKAs when compared to Col-0 and two different complemented lines. These results indicate that DIC2 is not involved directly in the transport of BCAAs or their derivatives.

Strikingly, all TCA cycle intermediates were significantly higher in abundance in *dic2-1* upon 10 days of dark treatment, whereas other mutants only displayed minor changes. To identify the TCA cycle metabolites that were affected the most by the absence of DIC2, we carried out a time-course measurement of organic acids contents over 12 days of prolonged darkness (Fig. 4C and Fig. S8C). When F_v_/F_m_ began to decline more rapidly in *dic2-1* on day 7 (Fig. S8A), only malate, 2-OG and citrate were significantly more accumulated. While all TCA cycle intermediates were significantly increased in the mutant 10 days after dark treatment, malate and 2-OG accumulated at least 10 times higher in abundance in the mutant than Col-0 and *dic2-1/gDIC2*. Citrate abundance in the Col-0 and *dic2-1/gDIC2* progressively declined from day 3 to day 12 of darkness, whereas it remained unchanged in *dic2-1* plants throughout the treatment. Notably, the accumulation of these metabolites by *dic2-1 in planta* would be consistent with a failure to properly regulate mitochondrial malate import and citrate export. Throughout a diurnal cycle, pool sizes of these metabolites were unchanged (Fig. 4A); it was only when plants were exposed to dark-induced starvation that altered patterns in the utilization of specific organic acids manifested (Fig. 4C).

### DIC2 modulates metabolic flux through TCA cycle and amino acid metabolism to support citrate export at night

We next further investigated the cause of the increased RN in *dic2-1*. An increased night-time consumption rate of sucrose, but not of glucose or fructose, by *dic2-1* was observed (Fig. 5A and Fig. S9A). The expression of several nutrient-responsive senescence markers, SAG101 (50), WRKY53 (51) and SEN1 (52), were also highly upregulated in the mutant in the dark but not in the light (Fig. 5B and Fig. S9B), suggesting night time starvation. These results, combined with a lack of photosynthetic differences (Fig. 3A, Fig. S6A-E), suggest that an accelerated depletion rate of carbon stores via respiration is leading to night time starvation in *dic2-1*. Thus, changes in metabolite abundances noted above were accompanied by a higher sucrose catabolism in the mutant at night. When leaf discs were incubated in the dark with uniformly labelled ^14^C-malate or ^14^C-leucine and the evolution of ^14^CO_2_ was monitored, we found that *dic2-1* showed higher ^14^CO_2_ emissions than Col-0 and *dic2-1/gDIC2* from malate but not leucine (Fig. 5C). The observed increase in BCAAs accumulation in *dic2-1* in the dark (Fig. 4A) was contributed by increased TCA cycle fluxes into biosynthetic pathways and/or an elevated proteolysis, while BCAAs breakdown for respiration remained unchanged. Thus, these data indicated that the loss of DIC2 results in increased sucrose utilization (or export for use by other tissues) and TCA cycle-facilitated respiration to compensate for a failure to maintain the homeostasis of organic acid oxidation.

**Figure 5.**
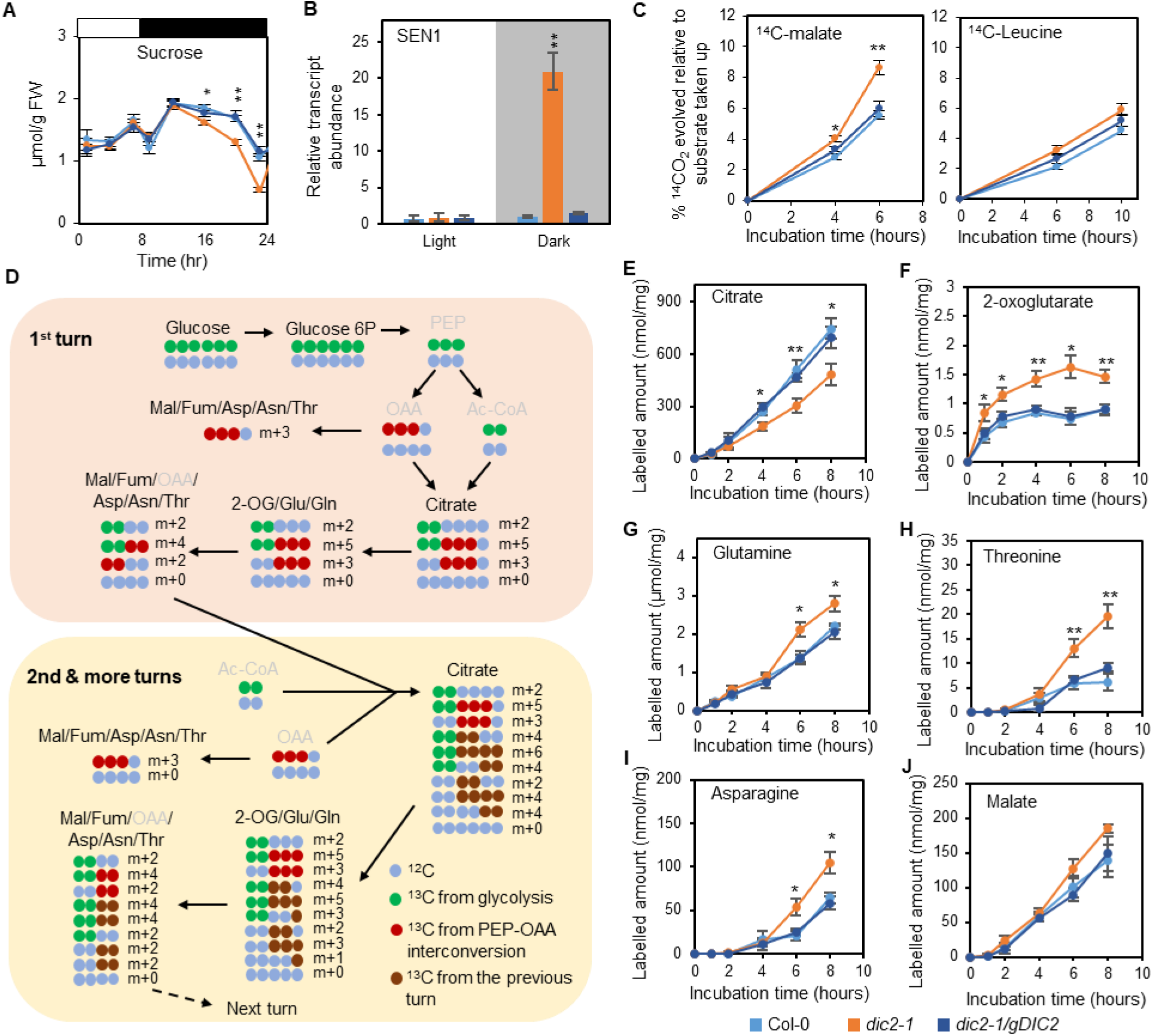
The loss of DIC2 causes an altered TCA cycle flux in the dark. (A) Sucrose levels in leaf discs from Col-0, *dic2-1* and *dic2-1/gDIC2* collected at different time points of a diurnal cycle (see Figure 3 legend) as quantitatively determined by GC-MS against authentic standards (n = 8). (B) qPCR analysis showing the expression of SEN1 in plants collected at the end of night (Shaded) or at the end of day (Light). All expression values were normalised against Col-0 end of day sample (n = 4). (C) CO_2_ evolution of leaf discs incubated in uniformly labelled ^14^C-malate (left) and ^14^C-leucine (right) in the dark. ^14^CO_2_ was captured in a NaOH trap, radiolabel in leaf discs was extracted and the radioactivity in these samples were counted by a liquid scintillation counter. Data shown is the percentage of CO_2_ released relative to the total amount of radiolabel incorporated into leaf metabolism (n = 3). (D) Schematic representation of all the possible incorporation patterns of isotope-labelled glucose into the TCA cycle via pyruvate dehydrogenase and/or phosphoenolpyruvate (PEP)-OAA interconversion in darkened leaf discs. Metabolites that were not measured are grey out. (E-F) Time-courses of ^13^C-labelling into metabolism in darkened leaf discs. Absolute abundance of total labelled citrate (E, sum of m+2, m+3, m+4, m+5 and m+6), 2-oxoglutarate (F, sum of m+2, m+3, m+4 and m+5), glutamine (G, sum of m+2, m+3, m+4 and m+5), threonine (H, sum of m+2 and m+4), asparagine (I, sum of m+2 and m+4) and malate (J, sum of m+2 and m+4) are shown. Means ± S.E. (n = 4). Asterisks indicating significant differences (* p < 0.05; ** p < 0.01) as determined by one-way ANOVA Tukey post-hoc analysis.

The observed decrease in substrate-dependent O_2_ consumption by isolated mitochondria (Fig. 2B) did not explain the faster dark respiration of intact leaves (Fig. 3E). This could be due to the absence of any extramitochondrial metabolism, which *in vivo* maintains metabolite supply for sustaining mitochondrial transport activities, TCA cycle and respiration in response to rearranged metabolism and transport in the mutant. To account for the apparent homeostasis in metabolite pool sizes (Fig. 4A), we postulated that there could be a flux change in certain steps of metabolism to compensate for the reduced malate import and citrate export from mitochondria. To determine if *dic2-1* metabolises carbon differently, we traced the flux of U-^13^C-glucose into the TCA cycle and closely related amino acids in leaf discs in the dark for 8 hours, and the ^13^C-tracing data was normalised by taking into consideration the differences in dark respiration rate (Fig. 5D; SI Dataset 1; SI Methods). Consistent with DIC2’s proposed function as a citrate transporter, *dic2-1* displayed a decreased rate of citrate labelling over the course of dark incubation (Fig. 5E). Such a decrease was unlikely to be contributed by a reduction in peroxisomal citrate synthase because labelled acetyl-CoA cannot be directly exported from mitochondria (53), and the existence of a carnitine/acylcarnitine carrier in plant mitochondria is questionable since its closest homolog BOU has recently been reported to transport glutamate (54). On the other hand, the citrate pool remained stable in the dark in *dic2-1* (Fig. 4A) possibly due to compensation by altered citrate turnover rates in other compartments. A decrease in OAA availability from cytosolic phosphoenolpyruvate (PEP) carboxylase could also be ruled out, since labelling of m+3 aspartate (a proxy for labelled OAA) was not altered (Fig. S10A). Decreased citrate labelling was accompanied by a significant increase in the abundance of labelled 2-OG due to a higher total pool in the mutant (Fig. 5F; SI Dataset 1), which coincided with an increased rate of ^13^C incorporation into glutamine (Fig. 5G). These increases could be facilitated by a higher mitochondrial glutamate efflux rate since there was a higher 2-OG accumulation in citrate-fed *dic2-1* mitochondria (Fig. S5E), implying that an enhanced flux into glutamine was necessary to remove excess mitochondrial 2-OG by mitochondrial glutamate transamination reactions in concert with plastidic glutamine synthase (55). Downstream extra-mitochondrial biosynthetic pathway of aspartate-derived amino acids, asparagine and threonine, were increased in abundances in the mutant (Fig. 5H-I), while the amount of labelled succinate, fumarate, malate and aspartate via the TCA cycle or PEP-OAA interconversion did not change (Fig. 5J and Fig. S10B-D). All these increases are consistent with a metabolic diversion of excess malate that was not consumed by mitochondria for citrate synthesis (due to a decrease in malate uptake and product inhibition of mMDH in the matrix) into aspartate. Overall, the ^13^C feeding data helped to explain how the mitochondrial phenotypes of *dic2-1* loss (Fig. 2) are significantly overcome in a whole plant metabolic context to establish day and night homeostasis and retain a viable plant albeit with stunted growth rate (Fig. 1). Only at day to night transitions and prolonged darkness do the consequences of these unusual metabolic fluxes yield temporary gross imbalances in metabolite pools.

## DISCUSSION

It was reported that DIC2, DIC1 and DIC3 are carriers for malate and OAA exchange to regulate NAD redox and photorespiration in the light (56–58). In addition, several mitochondrial malate carriers, including DIC2, were found to have a broad substrate preference for other dicarboxylates, tricarboxylates and/or amino acids with close resemblance to dicarboxylates *in vitro* (24–26), but their actual contributions to mitochondrial malate transport *in organello* or *in planta* have not been verified. Determining if these types of carriers are generalists or specialists *in vivo* is critical in any future attempt to modify mitochondrial substrate use, or to understand if transport is a point of control in metabolic models of plant cell function.

Malate import and citrate export are two of the most critical fluxes that mitochondria contribute to the rest of the cellular metabolic network in plants (59–61). Malate plays a pivotal role in mediating metabolic redox exchange between cellular compartments (57, 62). On the one hand, citrate produced in mitochondria is the major source of cellular citrate when the TCA cycle operates at night in mature green leaves. Here, we identified DIC2 as a single carrier linking these functions to mitochondrial anaplerotic metabolism. While our data reveal that other carriers can also mediate at least one of these two functions to provide partial functional backup, that is, however, insufficient to maintain metabolism unperturbed.

Given the prior evidence of broad DIC1 substrate preference *in vitro* and high amino acid identity among the DIC homologs, our finding that DIC1 could not complement the *dic2-1* phenotype (Fig. 1B, Fig. S2H) was unexpected (Fig. S11). Yet, inferring *in vivo* function from *in vitro* data on transport specificity data is known to be problematic and has prevented capture of the biological importance of transport of other substrates. For example, PAPST1 and PAPST2, which are 78% identical in amino acid sequence, both transport 3′-phosphoadenosine 5′-phosphate (PAP) and 3′-phosphoadenosine 5′-phosphosulfate (PAPS) in liposomes with only slight differences in their efficiency in driving ATP exchange (63). However, their true *in vivo* function was revealed only after additional genetic, biochemical and metabolomic approaches were employed: PAPST1 being responsible for the majority of PAPS/PAP transport to regulate glucosinolate biosynthesis in plastids, while PAPST2 fulfils a stress signalling role through PAP/AT(D)P exchange in chloroplasts and mitochondria (63, 64). In a similar way, while liposomes showed the ability of DIC2 to transport a variety of dicarboxylates and even citrate, albeit poorly (24), they could not accurately predict apparent specificity or directionality of transport in intact mitochondria (Fig. 2) or the *in vivo* consequences of DIC2 loss (Fig. 3-5).

Revealing the role for DIC2 in citrate export required moving to a less reductionist system, i.e. to the exploration of metabolic processes in isolated mitochondria (Fig. 2). Citrate produced in the mitochondria is predicted to be exported for fatty acid biosynthesis via the cytosolic ATP-dependent citrate lyase (65), or for storage in the vacuole based on flux prediction and ^13^C labelling analysis (20, 66), although it is also possible to fuel respiration in mitochondria isolated from storage tissues by externally supplying an excess amount of citrate (67). Given the importance of mitochondria-supplied citrate to metabolism in green tissues, it is expected that a complete block in mitochondrial citrate synthesis and export would result in severe reduction in post-germination vegetative growth. A null mutant of mitochondrial citrate synthase has not been reported to date, and a reduction in mitochondrial citrate synthase activity only resulted in a small change in vegetative growth phenotype (21, 22). Unlike in yeast and mammals, the function of plant mitochondrial pyruvate carrier (MPC) seems to be non-essential since its absence has no effect on growth or the cellular citrate pool (16, 68). In comparison, mutants lacking the activity in one of the subunits of mitochondrial pyruvate dehydrogenase complex (mPDC) or both mMDH isoforms, which catalyse acetyl-CoA and OAA formation respectively required for citrate synthesis, are significantly slower in vegetative growth (23, 69, 70). mPDC mutation causes a severe blockage in citrate generation in leaf (69) that does not appear to be compensated by altering pyruvate decarboxylation and citrate synthesis activities in other compartments. By contrast, OAA import into the mitochondria is sufficient to maintain a citrate pool in the mMDH double knockout similar to that in the wildtype at the expense of elevated respiration and increased accumulation of dicarboxylates (23). Here, we observed a reduction in vegetative growth in *dic2-1*, with decreases in citrate export by isolated mitochondria (Fig. 2F) and *in vivo* ^13^C-glucose labelling into citrate (Fig. 5E) when compared to the wildtype. Unlike mMDH and mPDC mutants, however, the steady state pool of organic acids, including citrate, was surprisingly stable in *dic2-1* (Fig. 4A). This suggests that fatty acid turnover is affected, either through increasing citrate synthesis via the last step of peroxisomal β-oxidation and/or decreasing citrate breakdown during the initial steps of plastidic fatty acid biosynthesis. The former is a more likely scenario given that *dic2-1* phenotypes are observed mostly during darkness when fatty acid biosynthesis is ineligible (71), and the transcripts of two peroxisomal citrate synthase isoforms are more abundant at night (72, 73) and during prolonged darkness (74). Lowering fatty acid accumulation by limiting triacylglycerol depletion helps plants survive prolonged darkness by dampening the degree of lipid peroxidation (75). Thus, the increased susceptibility to dark-induced senescence by *dic2-1* (Fig. 4B-C and Fig. S8) could be caused by a loss of synchronisation in fatty acid turnover with peroxisomal citrate synthesis and export, as a result of increased demand for peroxisomal citrate to compensate for a reduction in mitochondrial carbon supply. The requirement for an equilibrium between organellar citrate synthesis and fatty acid breakdown to prevent excess reactive oxygen species accumulation would reasonably explain why a strong DIC2 transcriptional response is commonly observed during abiotic stress treatments, including touch and sound vibration (76, 77), phosphate deficiency (78), light-to-dark transition (79) and cold, drought and UV stresses (80). Overall, these data strongly support a function of DIC2 in plant mitochondrial citrate transport, with a reduction in citrate export as a direct consequence of DIC2 loss, rather than a side effect.

The existence of multiple modes of TCA cycle substrate transport has been proposed (27, 28) but the lack of clarity as to the identity and substrate specificity of carriers responsible has been hindering our understanding on how they cooperate to support the integration of mitochondrial carbon metabolism into the wider cellular metabolic network (17). In the case of *dic2-1*, we found that there was no change in the uptake of externally supplied substrate by fully energised isolated mitochondria (Fig. S5 “INPUT”). Residual malate import and citrate export are maintained in *dic2-1* mitochondria (Fig. 2F, Fig. S5B), suggesting that other carriers exist to partially compensate for the absence of DIC2 activity. The majority of an added dicarboxylate was oxidised to pyruvate and/or OAA in mutant mitochondria with a diversion of excess malate destined for OAA and citrate formation into a hydration reaction to produce fumarate, which was then exported from isolated mitochondria (Fig. 2E) and redirected towards aspartate metabolism *in vivo* as seen by increases in asparagine and threonine labelling (Fig. 5H-I). It is remarkable that, despite all these changes, *in vivo* cytosolic NADH/NAD^+^ ratios in *dic2-1* leaves did not differ from wildtype throughout the night (Fig. 3D) with only a glimpse of change during a light-to-dark transition (Fig. 3C). We speculate that the homeostasis of NAD redox state in the mutant can be explained by at least one of the following: (i) an increased rate of malate oxidation being offset by a reduction in isocitrate oxidation in the cytosol; (ii) a rapid remobilisation of excess NADH into the intermembrane space of mitochondria where it was oxidised by external NADH dehydrogenase, thereby raising leaf respiration in the dark (Fig. 3E); and/or (iii) excess reductant used for diverting malate into threonine generation via aspartate catabolism, requiring 2 NADPH and 2 ATP molecules (81). Consistently, metabolite labelling and profiling have highlighted the flexibility of metabolism when multiple carriers with varying substrate preferences are present. For instance, the loss of mitochondrial citrate carrier in Drosophila larvae causes only a mild reduction in citrate content concomitant with increased malate and fumarate levels (82). Malate and citrate levels are slightly decreased in mice with suppressed expression of the mitochondrial dicarboxylate carrier (83). Silencing of the mitochondrial glutamate carrier in colorectal tumor cells resulted in a relatively small increase in cellular glutamate content whereas the aspartate level was reduced by ~45% (84). In plants, studies of mitochondrial carrier mutants reported to date are unable to directly link growth and/or metabolic phenotype with gene function (85–87). It is possible that, apart from functional redundancy, changes in metabolic fluxes occur without altering overall metabolite pool or physiology, which can only be captured by sensitive measurements under specific external conditions.

In conclusion, we have refined the current molecular identity of DIC2 to be a mitochondrial malate import and citrate export carrier in Arabidopsis, the absence of which leads to growth retardation. The importance of mitochondrial organic acid exchange carriers in plants has been discussed for decades, but it is only in the last decade that their identities are beginning to be unravelled through heterologous systems. By utilising a broad approach in combination with reverse genetics, however, we demonstrate that carrier analysis in heterologous systems are only the start to reveal the true nature of DIC2 *in vivo*. Given the lack of genetic clarity of many plant mitochondrial carriers, it is reasonable to suspect that the true identity of many other carriers has yet to be unravelled. Here, we provide a blueprint for characterising the *in organello* and *in vivo* function of a plant mitochondrial carrier that requires an integrated approach to analyse genetic lines. Such approaches reveal systems characteristics that may be strictly required from carrier function and specificity, such as exact membrane composition, bioenergetic status, pH and relative local metabolite pool sizes and fluxes. A complete, precise identification of mitochondrial metabolite carriers and their substrate preferences with kinetic characteristics and transport orientation will assist in improving genome scale metabolic models, as well as refining the role of DIC2 role in underpinning the remarkable flexibility of plant cell metabolism.

## METHODS

### Plant Material and Growth Conditions

The *DIC2* T-DNA insertion lines were obtained from GABI-Kat (*dic2-1* GK-217B05, *dic2-2* GK-833F11 and *dic2-3* GK-047F03)(88). gdh dKO (*gdh1-2 x gdh2-1*), *mcca1-1*, *mccb1-1* and *hml1-2* were obtained from Arabidopsis Biological Resource Center (CS860075, CS66518, CS66519 and CS66521 respectively (49, 89)). A detailed description of growth conditions, generation of transgenic lines and quantitative transcript analysis is provided in SI Methods.

### Isolation of mitochondria, O_2_ electrode measurements, enzyme activity assays and metabolite uptake assays by silicone oil centrifugation

Mitochondria were isolated from two-week-old Arabidopsis seedlings as described previously (90). O_2_ consumption by purified mitochondria was measured in a computer-controlled Clark-type O_2_ electrode unit according to Lee et al. (91). *In vitro* activities of TCA cycle enzymes in isolated mitochondria were measured as described by Huang et al. (92). Time-course measurements of substrate uptake by isolated mitochondria were carried out using silicone oil centrifugation technique according to Lee et al. (93) with modifications. Fractions above (extra-mitochondrial space) and bottom (pellet, mitochondria) were collected, and metabolites were methanol-extracted and detected by mass spectrometry. Additional information is provided in SI Methods.

### Analyses of metabolites by mass spectrometry

For GC-MS analysis of sugars, derivatised metabolite samples were analysed by an Agilent 6890 gas chromatograph coupled with a 7683B Automatic Liquid Sampler and a 5973N mass selective detector. For SRM-MS analysis of organic acids. For measuring organic acids amino acids, samples were analysed by an Agilent 1100 HPLC system coupled to an Agilent 6430 Triple Quadrupole (QQQ) mass spectrometer equipped with an electrospray ion source. A detailed description is provided in SI Methods.

### ^13^C-glucose labelling of Arabidopsis leaf discs and analysis of labelled metabolites

Leaf discs (~50 mg) were prepared from short-day grown (8-h light/16-h dark) plants 1 hour before the end of a normal light photoperiod. They were floated on leaf respiratory buffer containing 20 mM U-^13^C-glucose (99% purity, Sigma Aldrich). At the specified incubation time, leaf discs were briefly washed with respiratory buffer to remove excess labelled glucose and frozen in liquid nitrogen for metabolite extraction as stated above. Analyses of total, untargeted metabolites were performed using an Agilent 1100 HPLC system coupled to an Agilent 6510 Quadrupole/Time-of-Flight (Q-TOF) mass spectrometer equipped with an electrospray ion source. Peak extraction, isotopic correction for natural ^12^C abundances and analysis of isotopic abundances are described in SI Methods.

### Relative quantitation of mitochondrial protein abundances by LC-MRM-MS

Multiple reaction monitoring (MRM) was carried out exactly as described previously (94), except trypsin was added to the protein samples in a mass ratio of 1:20. Peptide abundances from each sample were normalised against VDAC in which its abundance was identical between mitochondria from Col-0, *dic2-1* and *dic2-1/gDIC2* based on western blotting. Transitions used for multiple reaction monitoring are provided in Table S3.

## Author Contributions

CPL performed most of the experiments in this manuscript and undertook data analysis. ME and PF conducted experiments with Peredox-mCherry lines and carried out data analysis. RF undertook MRM development and analysis. CPL, MS and AHM developed the experimental plan. CPL and AHM wrote the manuscript and all authors revised the manuscript.

## ACKNOWLEDGEMENT

This work was supported by the Australian Research Council (ARC) Centre of Excellence in Plant Energy Biology (A.H.M., grant number CE140100008) and the German Research Foundation (DFG) (M.S., grant number SCHW1719/5-1 as part of the package grant PAK918). We also thank Dr Brendan O’Leary (The University of Western Australia) for a critical reading of the manuscript.

