## Supplemental Information for "The Arabidopsis mitochondrial dicarboxylate carrier 2 maintains leaf metabolic homeostasis by uniting malate import and citrate export"

**This PDF file includes:**

Supplementary Methods

Figures S1 to S11

Tables S1 to S5

SI References

#### Supplementary Methods

##### Plant material and growth conditions

The *DIC2* T-DNA insertion lines were obtained from GABI-Kat (*dic2-1* GK-217B05, *dic2-2* GK-833F11 and *dic2-3* GK-047F03)(1). gdh dKO (*gdh1-2 x gdh2-1*), *dic2a1-1*, *dic2b1-1* and *hml1-2* were obtained from Arabidopsis Biological Resource Center (CS860075, CS66518, CS66519 and CS66521 respectively(2, 3)). These T-DNA insertion lines were screened for homozygous insertion by PCR of genomic DNA using the primer pairs according to Table S4. Inverse PCR was carried out according to Teschner et al. (4). The construct for genomic complementation with transgene gDIC2 was generated by amplifying the DIC2 genomic region (including ~2 kb upstream of start codon and ~1 kb downstream of stop codon) from wild type Col-0 DNA, and the resulting ~4 kb fragment was cloned into pCAMBIA1380. Overexpression constructs were generated by amplifying the coding sequence of DIC2, DIC1 or DIC3 and inserting each of these fragments into pB7WG2 by Gateway cloning. All cloning primers were listed in Table S4. Agrobacterium-mediated transformation of each construct was carried out by floral dipping.

Arabidopsis seedlings and plants were grown under a light intensity of 100 µmol $m^{-2}$ $s^{-1}$ after seeds were stratified at 4ºC in the dark for 2-3 days. Plants were grown on compost supplemented with Perlite and Vermiculite (3:1:1). For the isolation of mitochondria, seeds were surface-sterilized and grown in Murashige and Skoog (MS) medium (½-strength MS medium, 2 mM MES pH 5.7) supplemented with 1% (w/v) sucrose and 0.1% (w/v) agar for 14-16 days with gentle agitation (40-60 rpm) under long day conditions.

##### Subcellular localisation analysis by confocal laser scanning microscopy (CLSM)

A Nikon A1Si confocal microscope equipped with the following excitation and emission wavelength settings: 488-nm excitation and 525-nm emission for GFP, 560-nm excitation and 595-nm emission for TMRM and 640-nm excitation and 700-nm emission for chlorophyll autofluorescence. Images were acquired by a NIS element AR software package (version 4.13.01, Build 916) using a 20x lens (Nikon CFI Plan Apo VC 20x 0.75 N.A.) and a 60x lens (Nikon CFI Plan Apo VC 60x 1.20 N.A. WI) and were processed using ImageJ software package.

##### Transcript analysis by qRT-PCR

RNA isolation was performed using the Spectrum Plant Total RNA Kit (Sigma-Aldrich) in conjunction with on-column DNAase treatment to remove genomic DNA. cDNA was synthesised using the iScript cDNA Synthesis Kit (Bio-rad). Quantitative RT-PCR was performed in a LightCycler 480 II Real Time PCR System (Roche) using the QuantiNova

SYBR Green RT-PCR Kit (Qiagen) according to the manufacturer's instructions with reference genes *elongation factor-1α* (At5g60390) and *SAND family protein* (At2g28390)(5). All primers are listed in Table S4.

##### Isolation of mitochondria, $O_2$ electrode measurements, enzyme activity assays and metabolite uptake assays by silicone oil centrifugation

Mitochondria were isolated from two-week-old Arabidopsis seedlings as described previously (6). $O_2$ consumption by purified mitochondria was measured in a computer-controlled Clark-type $O_2$ electrode unit according to Lee et al. (7). *In vitro* activities of TCA cycle enzymes in isolated mitochondria were measured as described by Huang et al. (8).

Time-course measurements of substrate uptake by isolated mitochondria were carried out according to Lee et al. (9) with modifications. For uptake in the absence of cofactors, freshly prepared mitochondria (100 µg) were resuspended in 200 µL minimal transport buffer (225 mM sucrose, 1 mM $KH_2PO_4$, 10 mM TES, 0.5 mM ATP, pH 7.2). Uptake assays were initiated by adding radiolabeled U-[$^{14}$C]-malate (200 µM, Perkin Elmer) at 25°C. After the specified incubation time, the reaction was stopped immediately by rapid centrifugation (at 12,000 rpm for 1–2 min) through a 90 µL silicone oil layer (AR200, Sigma Aldrich) into the bottom sedimentation layer containing 20 µL of 10% (v/v) perchloric acid. Fractions above (supernatant) and below (containing the mitochondria pellet) the silicone oil interface were transferred to separate scintillation vials and mixed with 3 ml of Ultima Gold™ scintillation fluid (PerkinElmer). Radioactivity was detected by a Beckman LS 6500 Scintillation Counter. For energised uptake, isolated mitochondria (100 µg) were resuspended in 190 µL respiratory transport medium (225 mM sucrose, 5 mM $KH_2PO_4$, 10 mM TES, 10 mM NaCl, 4 mM $MgSO_4$, 0.1% (w/v) BSA, 2 mM NAD, 12 µM CoA, 0.2 mM thiamine pyrophosphate and 0.5 mM ADP, pH 7.2) and uptake was initiated by adding 50 µM or 500 µM substrate as stated. Reactions were stopped as described above, except the bottom layer contained 20 µL of 0.5 M sucrose pH 1.5. Fractions above (extra-mitochondrial space) and bottom (pellet, mitochondria) were collected, and metabolites were methanol-extracted and detected by LC-SRM-MS (see below).

##### Blue-native gel electrophoresis, SDS gel electrophoresis and immunoblotting

Separation of digitonin-solubilized mitochondrial proteins was carried out according to Eubel et al.(10), using a 4.5%-16% gradient BN gel dimension. In-gel Complex I activity staining was carried out as outlined by Sabar et al. (11).

##### Leaf gas exchange, chlorophyll fluorescence and respiration measurements

Measurements of gas-exchange parameters were carried out using a LI-6400 XT infrared gas analyser (Li-Cor). After at least two hours of illumination, measurements were carried out using fully developed leaves from at least 8-week-old short-day grown plants. Prior to each measurement, leaf was acclimated in the enclosed 60mm$^2$ chamber to 22°C, relative humidity 70% and a $CO_2$ concentration of 400 ppm with light intensity of 250 µmol m$^{-2}$ s$^{-1}$. A series of light intensities (0, 50, 100, 250, 500, 700 µmol m$^{-2}$ s$^{-1}$) were applied and the following parameters were then recorded: $CO_2$ assimilation rate, stomatal conductance and transpiration rate. For the analysis of post-illumination bursts, leaf was equilibrated in a chamber to 22°C and relative humidity 70%, with light intensity of 1000 µmol m$^{-2}$ s$^{-1}$ and $CO_2$ concentration of 100 ppm. $CO_2$ assimilation rate in the light was monitored for 3 min before dark shift (i.e. PAR = 0 µmol m$^{-2}$ s$^{-1}$) and $CO_2$ evolution rate was monitored for 2.5 minutes.

Chlorophyll fluorescence measurements were carried out using the IMAGING-PAM Maxi Version Chlorophyll Fluorometer (WALZ). Plant was acclimated in the dark for at least 15 minutes before $F_v/F_m$, PSII quantum yield during illumination (Y(II)), photosynthetic electron transport rate (ETR) and non-photochemical quenching (NPQ) were monitored at increasing actinic light intensities and plotted as light response curves.

Dark respiration was measured in dark-adapted, fully expanded leaves using a Q2 $O_2$ sensor (Astec-Global) as previously described (12). $O_2$ concentration in a sealed 850 µL capacity tube with a single leaf disc (6.5 mm diameter, in a total of 40-60 mg fresh weight) floated on leaf respiration buffer (10 mM HEPES, 10 mM MES and 0.2 mM $CaCl_2.2H_2O$, pH 7.2) was measured.

**Analysis of NAD redox dynamics**

All experiments were carried out with leaf discs of 6-week-old *Arabidopsis thaliana* plants grown under long-day conditions (16h light/ 8h dark). Microscopy procedures described in Wagner et al. (13) were used as guideline for ratiometric sensor monitoring with the modification that samples were mounted between two cover slips (22 x 40 mm, VWR).. Dynamic changes under dark-light conditions were monitored with a confocal laser scanning microscope (Zeiss LSM 780, attached to an Axio Observer.Z1; Carl Zeiss Microscopy) using the 10x objective (Plan-Apochromat, 0.3 M27). Excitation was set at 405 nm (T-Sapphire) and 543 nm (mCherry), emission was collected at 499–544 nm (T-Sapphire) and 588–624 nm (mCherry). The ZEN Experiment Designer software module (Carl Zeiss) was used to program time series with varying scanning cycles and illumination times. A custom made device with an implemented cold white LED stripe (Pferdekaemper, Menden, Germany) was used to apply actinic light with an intensity of 220 µmol m$^{-2}$ s$^{-1}$.Time series with Peredox-mCherry were performed with the following regime: (i) 15 min in the dark of 30 scans with 30 seconds interval;

(ii) 15 min of illumination with 8 loops of 15 seconds light followed by 3.2 seconds scan and 26 loops of 30 seconds light followed by 3.2 seconds scan; (iii) 15 min in the dark of 30 scans with 30 seconds interval.

For *in vivo* measurements of NAD redox changes, leaf discs stably expressing cytosolic Peredox-mCherry were subjected to multiwell plate reader-based fluorimetry, as described in De Col et al. (2017). Plant material was submerged in 200 µL assay medium (10 mM HEPES, 10 mM MES and 0.2 mM $CaCl_2$, pH 7.2) in a transparent 96-well plate (NUNC). Peredox-mCherry fluorescence was recorded in a CLARIOstar microplate reader. The chromophores T-Sapphire and mCherry were excited at 400 ± 10 nm and 570 ± 10 nm, and emission collected at 515 ± 7.5 nm and 610 ± 5 nm, respectively. Samples were imaged employing top optics with focal height of 8.0 mm, incubation temperature at 25°C, well-multichromatic monitoring, 50 flashes per cycle and double orbital shaking at 400 rpm for 10 s before each measurement cycle. Background fluorescence of control plants without sensor expression was recorded in parallel and subtracted from all data before analysis. Mounting of the leaf discs was performed, except for the use of a green LED head torch, in the dark at the end of the night shortly prior night-to-day transition. After recording the baseline, plates were exposed to actinic light under white LEDs with a photon flux density of 120 µmol $m^{-2}$ $s^{-1}$ for in total 8 h, except for 3-min-long interruptions allowing for fluorometric sensor recording at indicated time points. For the subsequent dark phase, plate was kept inside the plate reader (dark) and measurements were performed every 30 min.

**Metabolite Extraction**

Plant tissues (~25 mg for general metabolite analysis or ~50 mg for isotope labelling experiments) were collected at specified time points and immediately snap-frozen in liquid nitrogen. Samples were ground to fine powder and 500 µL of cold metabolite extraction solution (90% [v/v] methanol, spiked with 2 mg/mL ribitol, 6 mg/ml adipic acid, and 2 mg/ml and $^{13}$C-leucine as internal standards). Samples were immediately vortexed and shaken at 1,400 rpm for 20 min at 75°C. Cell debris was removed by centrifugation at 20,000 x g for 5 minutes. For each sample, 100 µL (400 µL for isotope labelling experiments) of supernatant was transferred to a new tube and either proceeded to derivatization for LC-MS analysis or dried using a SpeedVac.

**Analyses of sugars by gas chromatography mass spectrometry (GC-MS)**

For GC-MS analysis of sugars, dried samples were dissolved in 20 µL of pyridine in methoxylamine hydrochloride (20 mg $mL^{-1}$) and incubated at 30°C for 90 min at 1200 rpm. Then, 30 µL of N-methyl-N-(trimethylsilyl)-trifluoroacetamide was added to each sample and allowed to react for 30 min at 37°C at 1,400 rpm. The derivatised metabolite samples were analysed by an Agilent 6890 gas chromatograph coupled with a 7683B Automatic Liquid Sampler and a 5973N mass selective detector, and fitted with an Agilent VF-5ms capillary column (30 m x 0.25 μm, 0.25 mm internal diameter) and a 10 m integrated guard column. The helium carrier gas flow rate was 1 mL/min. For the 58.5-minute temperature gradient, the GC oven was held at the initial temperature of 70°C for 1 min, increased to 76°C at the rate of 1°C per minute and then to 325°C at the rate of 6°C per minute where it was held at 325°C for 10 min. The inlet temperature, MS source temperature and quadrupole temperature were 300°C, 230°C and 150°C respectively. The MS detector m/z scan range was 40 to 600.

**Analyses of organic acids and amino acids by multiple reaction monitoring using liquid chromatography selective reaction monitoring mass spectrometry (LC-SRM-MS)**

For LC-MS analysis of organic acids, sample derivatization was carried out based on previously published method with modifications (14). Briefly, for each of 100 µL of sample, 50 µL of 250 mM 3-nitrophenylhydrazine in 50% methanol, 50 µL of 150 mM 1-ethyl-3-(3-dimethylaminopropyl) carbodiimide in methanol, and 50 µL of 7.5% pyridine in 75% methanol were mixed and allowed to react on ice for 60 minutes. To terminate the reaction, 50 µL of 2 mg $mL^{-1}$ butylated-hydroxytoluene in methanol was added, followed by the addition of 700 µL of water. Derivatized organic acids were separated on a Phenomenex Kinetex XB-C18 column (50 x 2.1mm, 5µm particle size) using 0.1% formic acid in water (solvent A) and methanol with 0.1% formic acid (solvent B) as the mobile phase. The elution gradient was 18% B at 1 min, 90% B at 10 min, 100% B at 11 min, 100% B at 12 min, 18% B at 13 min and 18% B at 20 min. The column flow rate was 0.3 mL/min, and the column temperature was maintained at 40 °C. The QQQ-MS was operated in negative ion mode in selective reaction monitoring (SRM) mode.

For measuring amino acids, dried samples were resuspended in 500 µL HPLC-grade water before they were filtered to remove insoluble debris. Metabolites were separated on an Agilent Poroshell 120 Bonus-BP column (100 x 2.1 mm, 2.7µm internal diameter) using 0.1% formic acid in water (solvent A) and acetonitrile with 0.1% formic acid (solvent B) as the mobile phase. For the analysis of amino acids and sugars, the elution gradient was 0% B at 1 min, 1% B at 4 min, 10% B at 6 min, 100% B at 6.5 min, 100% B at 8 min, 0% B at 8.5 min and 0% B at 15 min. The column flow rate was 0.25 mL/min, the column temperature was kept at 40°C. The QQQ-MS was operated in positive ion mode in SRM mode.

A 0.5 µL or a 15 µL aliquot of each sample were injected and analysed by an Agilent 1100 HPLC system coupled to an Agilent 6430 Triple Quadrupole (QQQ) mass spectrometer equipped with an electrospray ion source. Data acquisition and LC-MS control were done using the Agilent MassHunter Data Acquisition software (version B06.00 Build 6.0.6025.4).

The autosampler was kept at 10°C. The QQQ-MS was operated in SRM mode using the following operation settings: capillary voltage, 4000V; drying N2 gas and temperature, 11 L/min and 125 °C respectively; Nebulizer, 15 psi. All optimised SRM transitions for each target were listed in Table S5. Data analysis was carried out using MassHunter Quantitative Analysis Software (version B.07.01, Build 7.1.524.0). Metabolites were quantified by comparing the integrated peak area with a calibration curve obtained using authentic standards, and normalised against fresh weight and internal standards.

**Dynamic $^{13}$C-glucose labelling of Arabidopsis leaf discs and analysis of labelled metabolites by time-of-flight mass spectrometry (TOF-MS)**

Leaf discs (~50 mg) were prepared from short-day grown (8-h light/16-h dark) plants 1 hour before the end of a normal light photoperiod. They were floated on leaf respiratory buffer containing 20 mM U-$^{13}$C-glucose (99% purity, Sigma Aldrich). At the specified incubation time, leaf discs were briefly washed with respiratory buffer to remove excess labelled glucose and frozen in liquid nitrogen for metabolite extraction as stated above.

Analyses of unlabelled and labelled metabolites were performed using an Agilent 1100 HPLC system coupled to an Agilent 6510 Quadrupole/Time-of-Flight (Q-TOF) mass spectrometer equipped with an electrospray ion source. Data acquisition and LC-MS control were carried out using the Agilent MassHunter Data Acquisition software (version B.02.00). Separation of metabolites was performed using a Luna C18 column (Phenomenex; 150 × 2 mm, 3 µm particle size). The mobile phase consisted of 97:3 water:methanol with 10 mM tributylamine and 15 mM acetic acid (solvent A) and 100% methanol (solvent B). The gradient program was 0% B 0 min, 1% B 5 min, 5% B 15 min, 10% B 22 min, 25% B 24 min, 27% B 35 min, 29% B 80 min, 95% B 81 min, 95% B 82 min, 0% B 83 min and 0% B 97 min. The flow rate was 0.2 mL/min, with column temperature kept at 35°C, autosampler was cooled to 10°C and injection volume was 30 µL. The Q-TOF was operated in MS mode with negative ion polarity using the following operation settings: capillary voltage, 4000V; drying N2 gas and temperature, 10 L/min and 250 °C respectively; Nebulizer, 30 psi. Fragmentor, skimmer and octopole radio frequency (Oct1 RF Vpp) voltages were set to 110V, 65V and 750V respectively. The scan range was 70-1200 m/z and spectra were collected at 4.4 spectra/s which corresponded to 2148 transients/spectrum. MS scan peaks of all the possible mass isotopologues were integrated using MassHunter Quantitative Analysis Software (version B.07.01, Build 7.1.524.0). Extracted peak matrices were processed using IsoCor (15) to correct for the contribution of naturally occurring isotopes. Relative isotopomer abundance for each metabolite and percentages of $^{13}$C enrichment were calculated according to Araújo et al. (16). Metabolite quantitation was carried out based on calibration curves obtained with unlabelled authentic standards. Incorporation values for *dic2-1* are corrected by dividing these numbers by a factor of 1.3, which was the fold increase in dark respiration for mutant leaf discs when compared to the wildtype (Fig. 2E).

#### Supplementary Figures

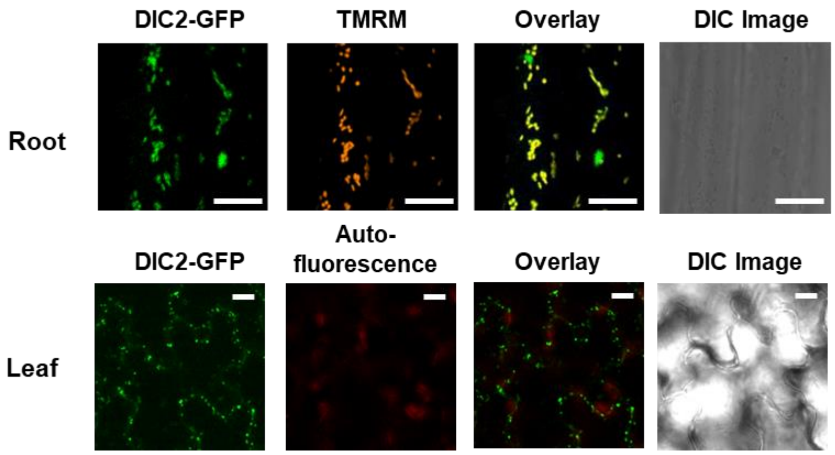


**Figure S1. Subcellular localization of MCC in Arabidopsis.** (A) Confocal laser scanning microscopy image of root cells (upper panels) and leaf epidermis from transgenic plants stably expressing MCC-GFP under the control of a UBQ10 promoter. GFP signal is pseudocolored green, tetramethylrhodamine (TMRM) is pseudocolored orange and chlorophyll fluorescence is pseudocolored red. Scale bar = 10 µm.

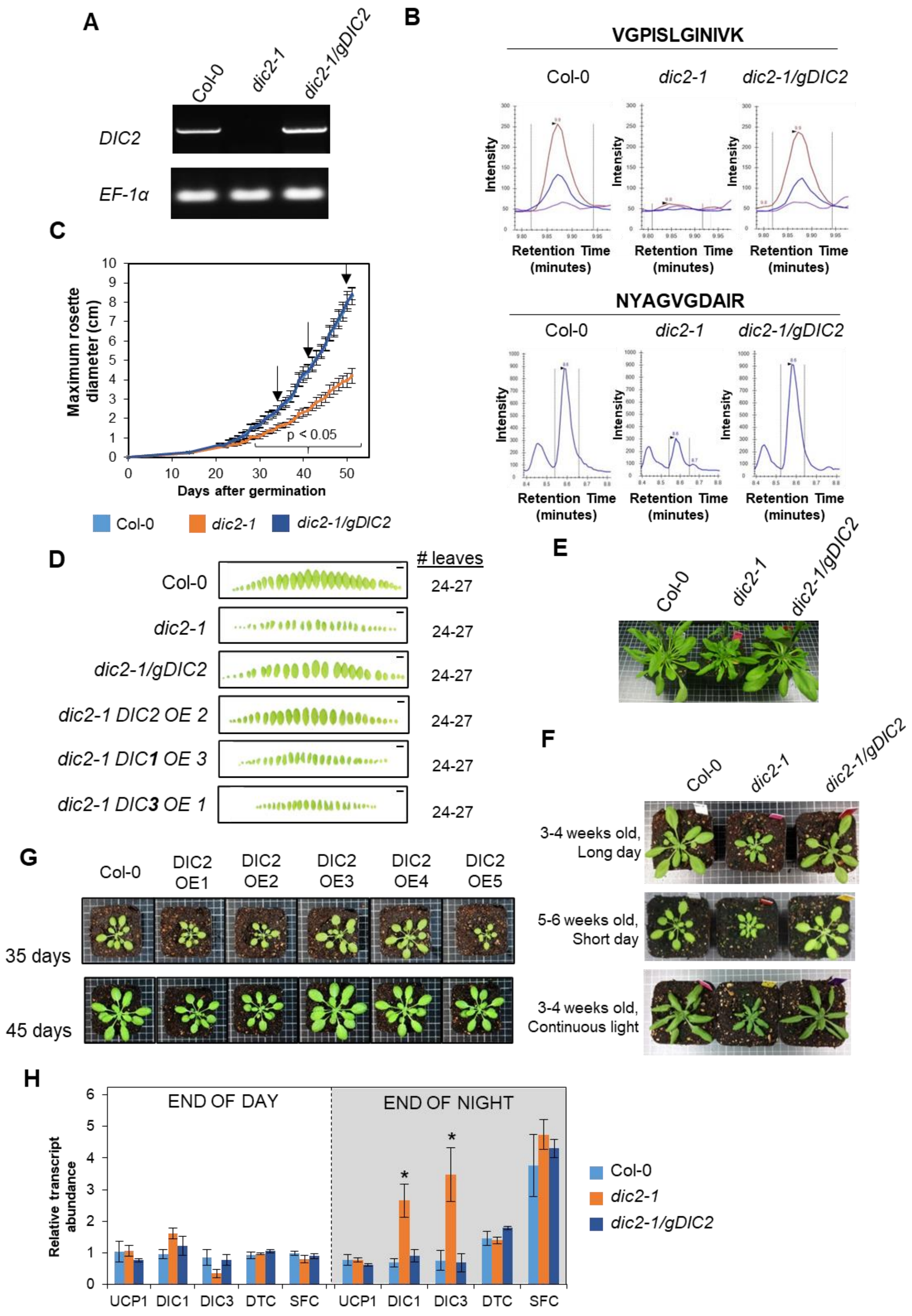


**Figure S2. Phenotypes of *dic2-1* and *dic2-1/gDIC2* plants.** (A) Semi-quantitative RT-PCR with primers amplifying full length DIC2, with elongation factor 1 alpha subunit (EF-1α) as a loading control. (B) LC-SRM-MS analysis showing representative chromatographic profiles for VGPISLGINIVK and NYAGVGDAIR peptides in mitochondria isolated from Col-0, *dic2-1* and *dic2-1/gDIC2*. Coloured lines correspond to different product ions generated from the fragmentation of

VGPISLGINIVK peptide. Integrated area of the highest peak (quantifier) from different genotypes is compared in Figure 1. (C) Comparison of rosette expansion rate by measuring rosette diameter over 49 days of growth. Arrows correspond to 35, 42 and 49 days after germination as shown in Figure 1B (n = 9). Significant differences ($p < 0.05$) were detected from Day 30 (between Col-0 vs *mcc-1* and *mcc-1* vs *mcc-1/gMCC* pairs based on one-way ANOVA Tukey post-hoc test) (D) Leaf numbers and phenotypes of different genotypes 42 days after germination. The range of leaf numbers from 5 plants is given on the right. Scale bar = 1 cm. (E) Vegetative phenotype one week after the first inflorescence was visible. (F) Phenotype of Col-0, *dic2-1* and *dic2-1/gDIC2* in ~3-4 weeks of long-day, ~5-6 weeks of short-day and 3-4 weeks of continuous light growth conditions. (G) Phenotype of Col-0 and DIC2 overexpression lines grown on soil under short day condition (8 h light/16 h dark) on 35, 45 days after germination. Representative top view of various genotypes is shown. Expression levels are shown in Table S1. (H) qPCR determination of relative transcript abundances of UCP1, DIC1, DIC3, DTC and SFC in the indicated genotypes collected at the end of day and the end of night. All expression values were normalised against Col-0 end of day sample (n = 4). Values shown are mean ± standard error, with asterisks (*) indicating significant differences ($p < 0.05$) between Col-0 vs *mcc-1* and *mcc-1* vs *mcc-1/gMCC* pairs as determined by one-way ANOVA Tukey post-hoc test.

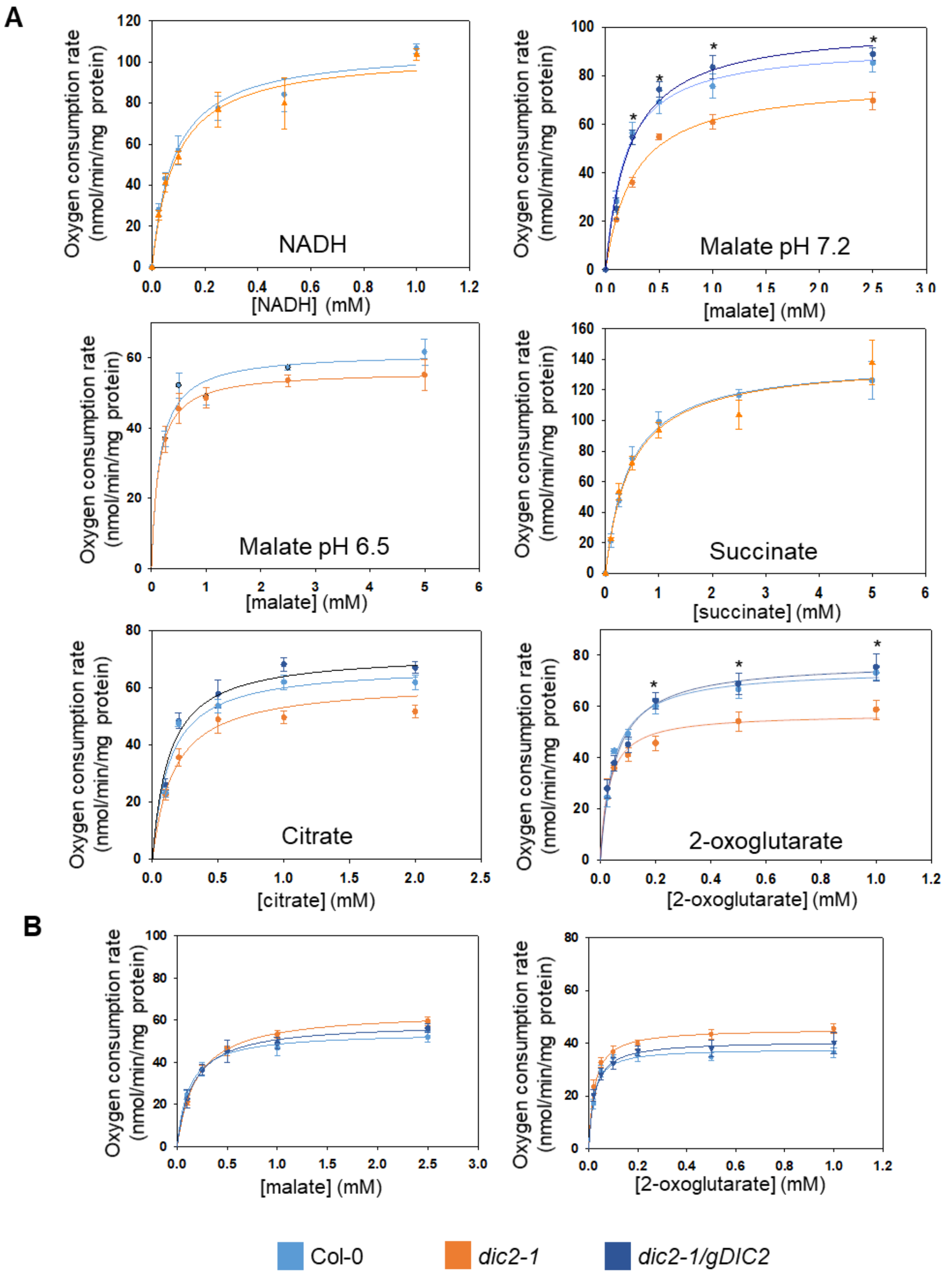


**Figure S3. Respiratory capacity of mitochondria isolated from the three genotypes.** (A) State III respiration of 100 µg mitochondrial proteins was assayed at varying concentration of NADH, succinate, malate at pH7.2 and 6.5, 2-OG and citrate (0.1 – 5 mM) using a Clarke-type $O_2$ electrode. (B) State IV respiration of 100 µg mitochondrial proteins was assayed at varying concentration of malate (left) and 2-OG (right) (0.1 – 2.5 mM) using a Clarke-type $O_2$ electrode. Values are mean ± S.E. (n = 5). * $p < 0.05$ from one-way ANOVA with Tukey post-hoc test.

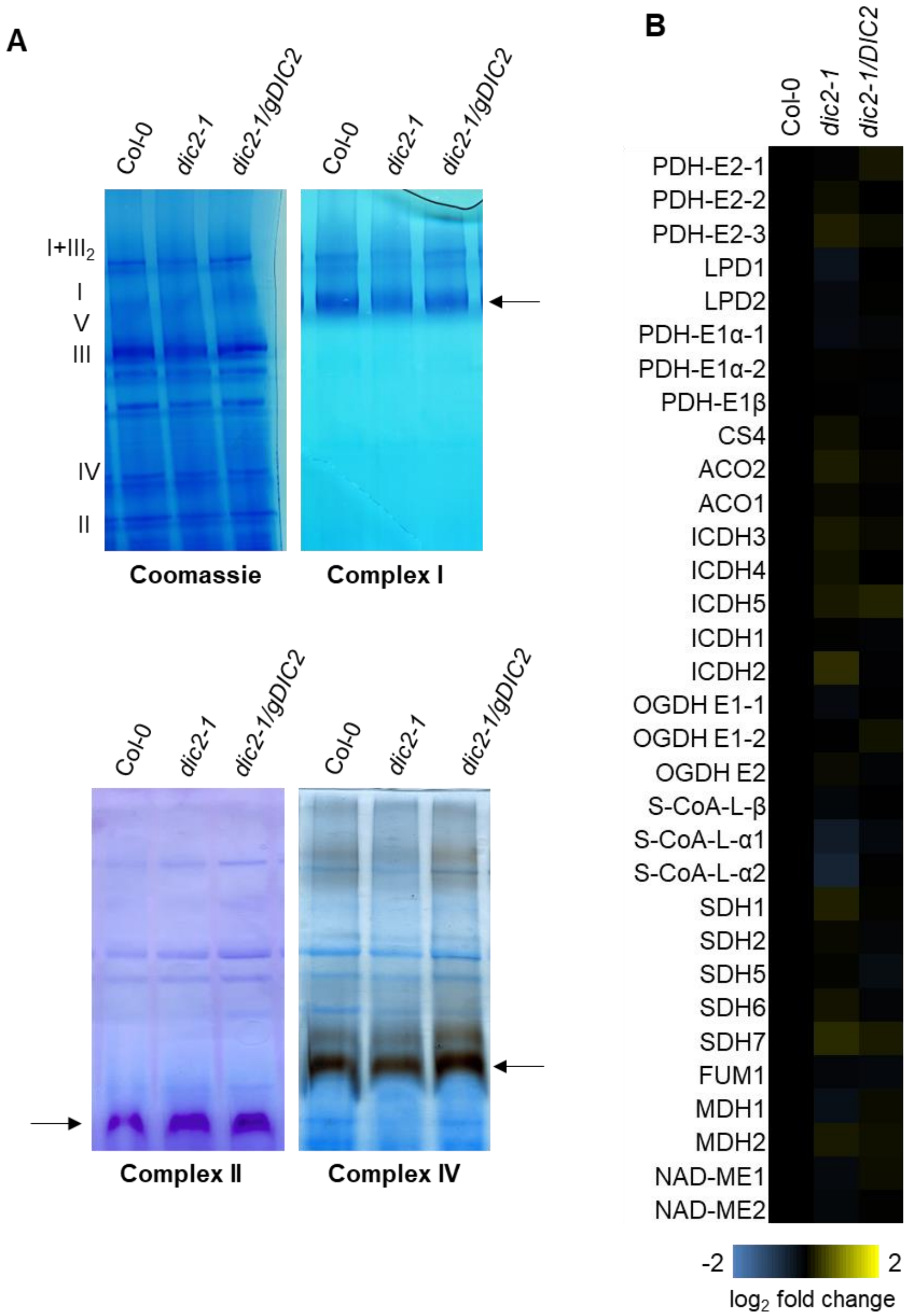


**Figure S4. An extensive survey of substrate feeding to isolated mitochondria.** (A) Separation of mitochondrial respiratory supercomplexes by 1D-blue-native PAGE. Roman numerals correspond to the locations of respiratory complexes. Gels were visualised by Coomassie Blue, Complex I, Complex II or Complex IV staining. (B) Heat map showing $\log_2$-fold change in the abundance of proteins in the TCA cycle and related pathways, relative to Col-0, as determined by LC-MRM-MS ($n \geq 3$). Abbreviations: PDH-E1 pyruvate dehydrogenase E1 subunit; PDH-E2, pyruvate dehydrogenase E2 subunit; LPD, lipoamide dehydrogenase (PDH-E3); CS, citrate synthase; ACO, aconitase, ICDH, isocitrate dehydrogenase; OGDH-E1, 2-oxoglutarate dehydrogenase E1 subunit; OGDH-E2, 2-oxoglutarate dehydrogenase E2 subunit; S-CoA-L, succinyl-CoA ligase; SDH, succinate dehydrogenase; FUM, fumarase; MDH, malate dehydrogenase; NAD-ME, NAD-dependent malic enzyme.

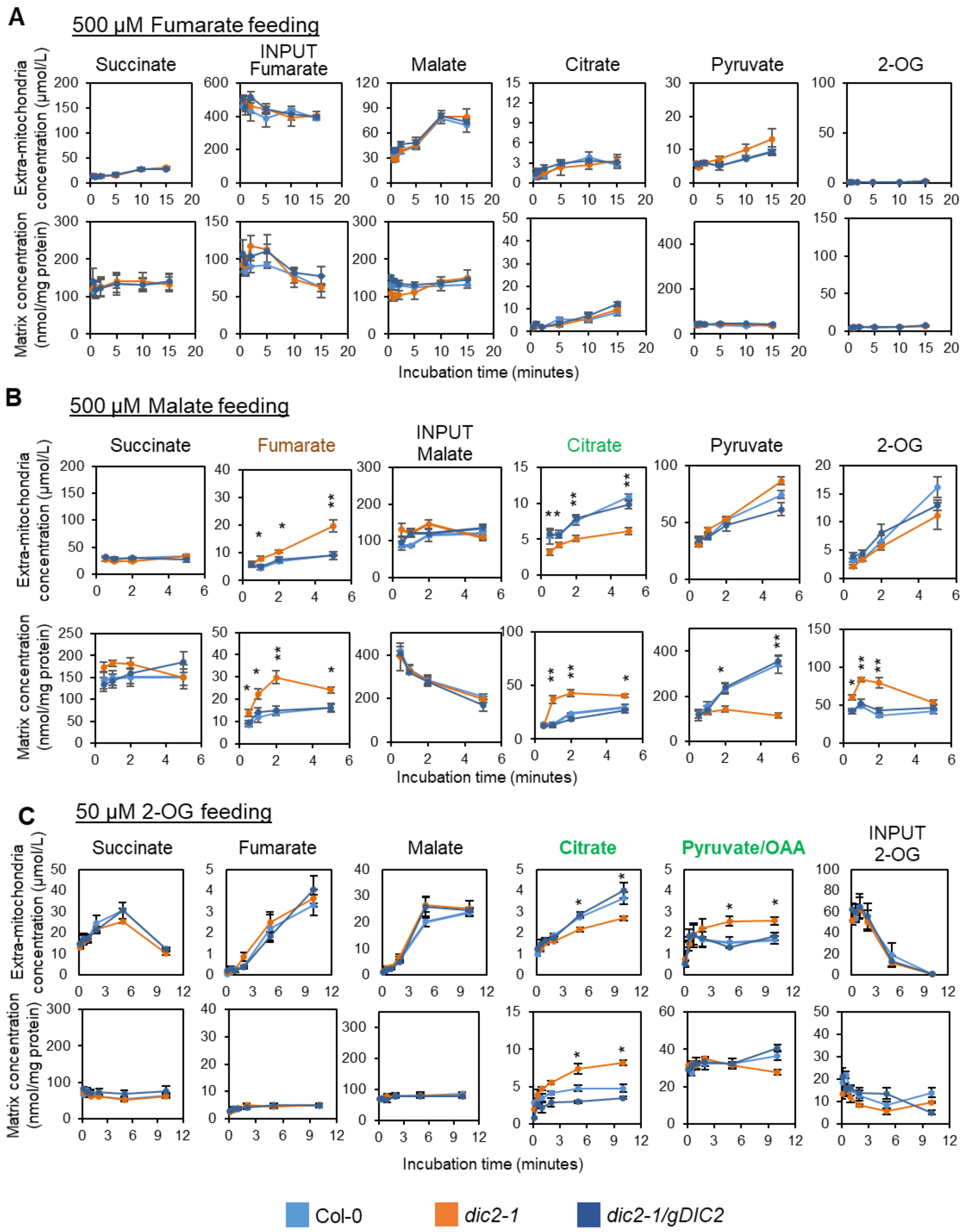


**Figure S5. (continue on the next page)**

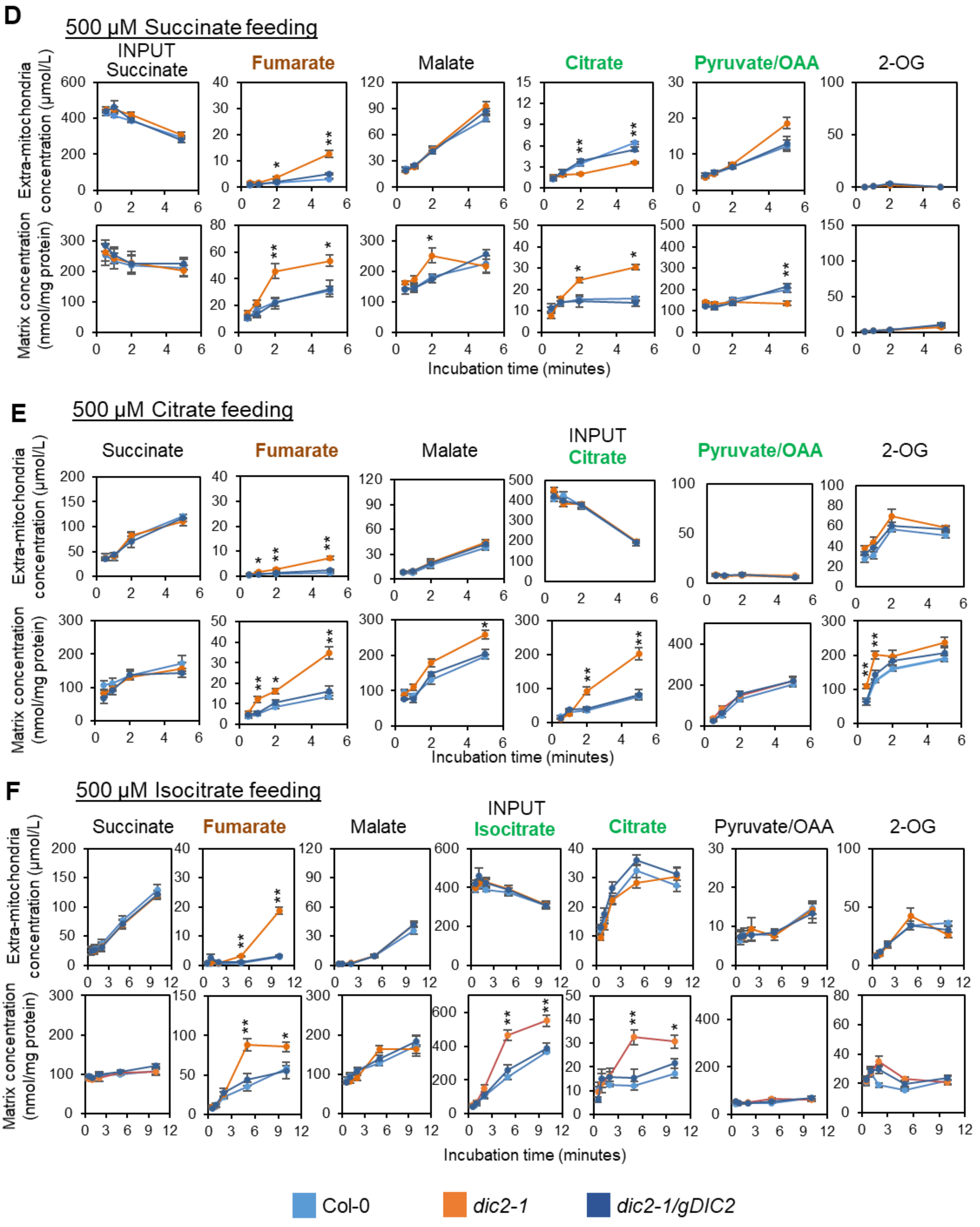


**Figure S5. An extensive survey of substrate feeding to isolated mitochondria.** Time-courses of metabolite concentrations in the extra-mitochondrial space (upper panels) and matrix (lower panels) incubated with different substrates and concentrations. Isolated mitochondria were incubated with substrates together with cofactors to stimulate state III respiration. Metabolic reaction was stopped by centrifugation in which the mitochondrial pellet (to measure matrix concentration) was separated from supernatant containing the substrate and exported products (to measure extra mitochondria metabolite concentrations) by a single silicone oil layer. Each data point represents mean ± S.E., n = 4. Asterisks denote significant differences between *dic2-1* vs Col-0 and *dic2-1* vs *dic2-1/gDIC2* based on ANOVA and Tukey's post-hoc analysis (* $p < 0.05$; ** $p < 0.01$).

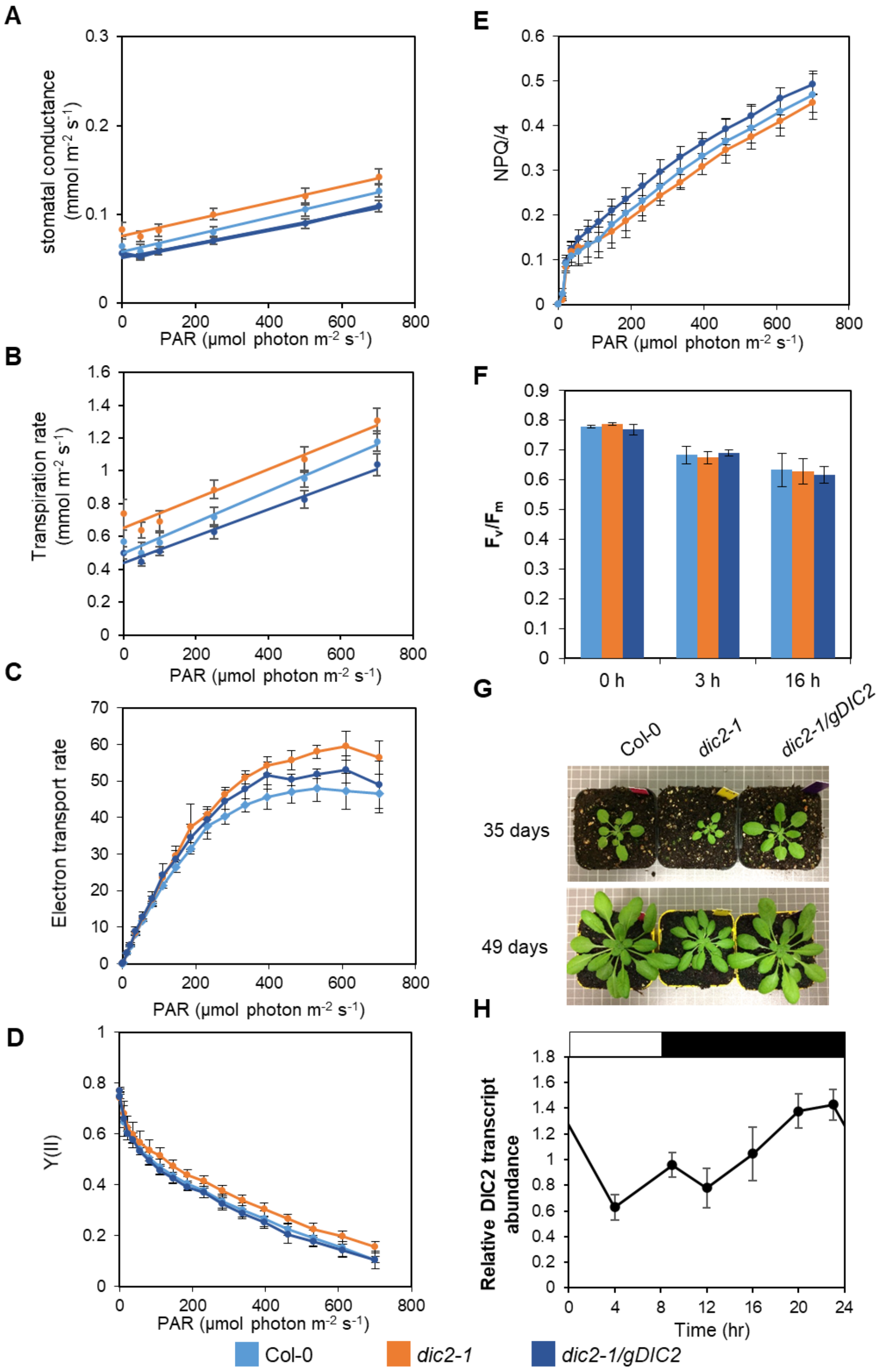


**Figure S6. Changes in photosynthetic and respiratory parameters in the DIC2 knockout mutant.** (A and B) Stomatal conductance (A) and transpiration rate (B) at different photosynthetic active radiation (PAR) with $CO_2$ concentration at 400 p.p.m. and temperature at 22°C (n = 6). (C-F) Light response curves of relative photosystem II electron transport rate (C), quantum efficiency of photosystem II (Y(II), D) and non-photochemical quenching (NPQ; E) of plants dark-adapted for at least 15 minutes (n = 6). (F) The photochemical efficiency of photosystem II ($F_v/F_m$) of plants treated with 3- or 16-hour high light stress (500 µmol $m^{-2}$ $s^{-1}$; n = 5). Each data point represents means ± SE. No significant differences were found based on one-way ANOVA analyses. (G) Phenotype of Col-0, *dic2-1* and *dic2-1/gDIC2* grown under short day conditions in 0.2% $CO_2$ (non-photorespiratory condition) after 35 days and 49 days of germination. (H) Relative expression of DIC2 in a diurnal cycle. Data represents average ± S.E. (n = 4).

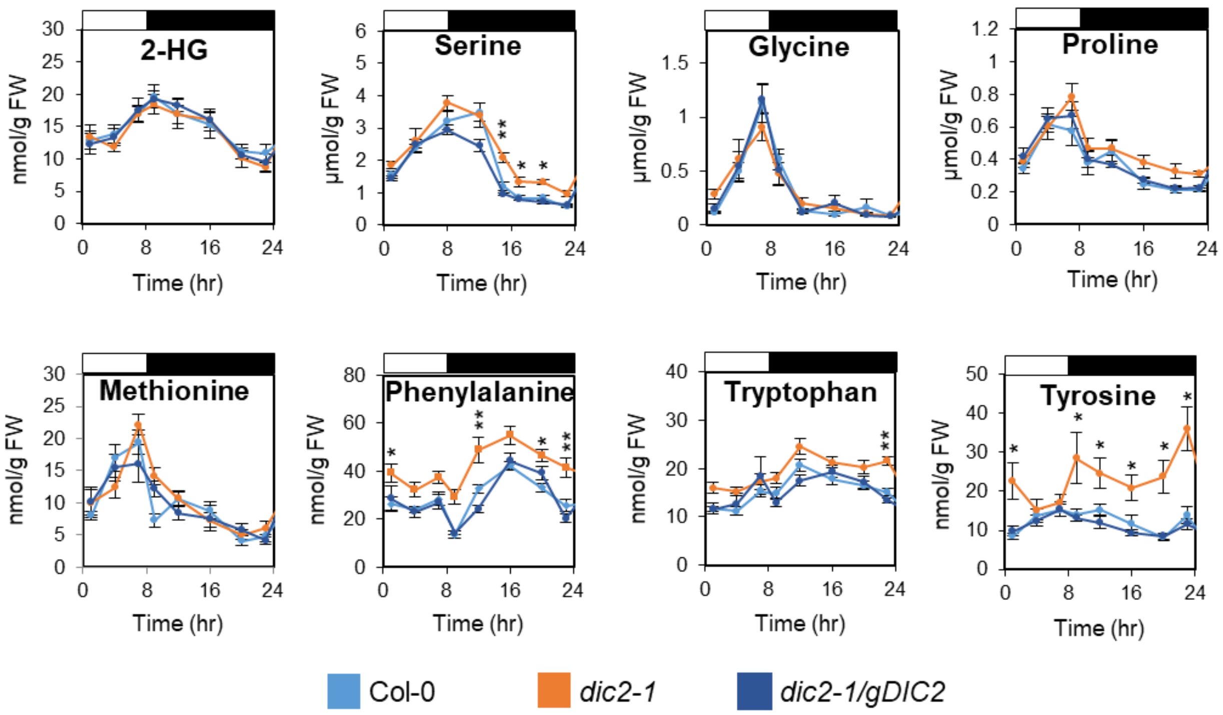


**Figure S7. Profile of selected amino acids in a diurnal cycle.** Plants were grown under short day conditions for 6 weeks and leaf discs were collected at 1, 4, 8, 12 and 15 hours after dark shift and 1, 4 and 7 hours after light shift. All the metabolites in this figure were analysed by GC-MS and LC-SRM-MS. Each data point represents mean ± S.E. ($n \geq 6$) in either nmol or µmol per gram fresh weight (FW), asterisks indicate a significant change as determined by the one-way ANOVA with Tukey post-hoc test (* $p < 0.05$; ** $p < 0.01$). Abbreviation: 2-HG, 2-hydroxyglutarate.

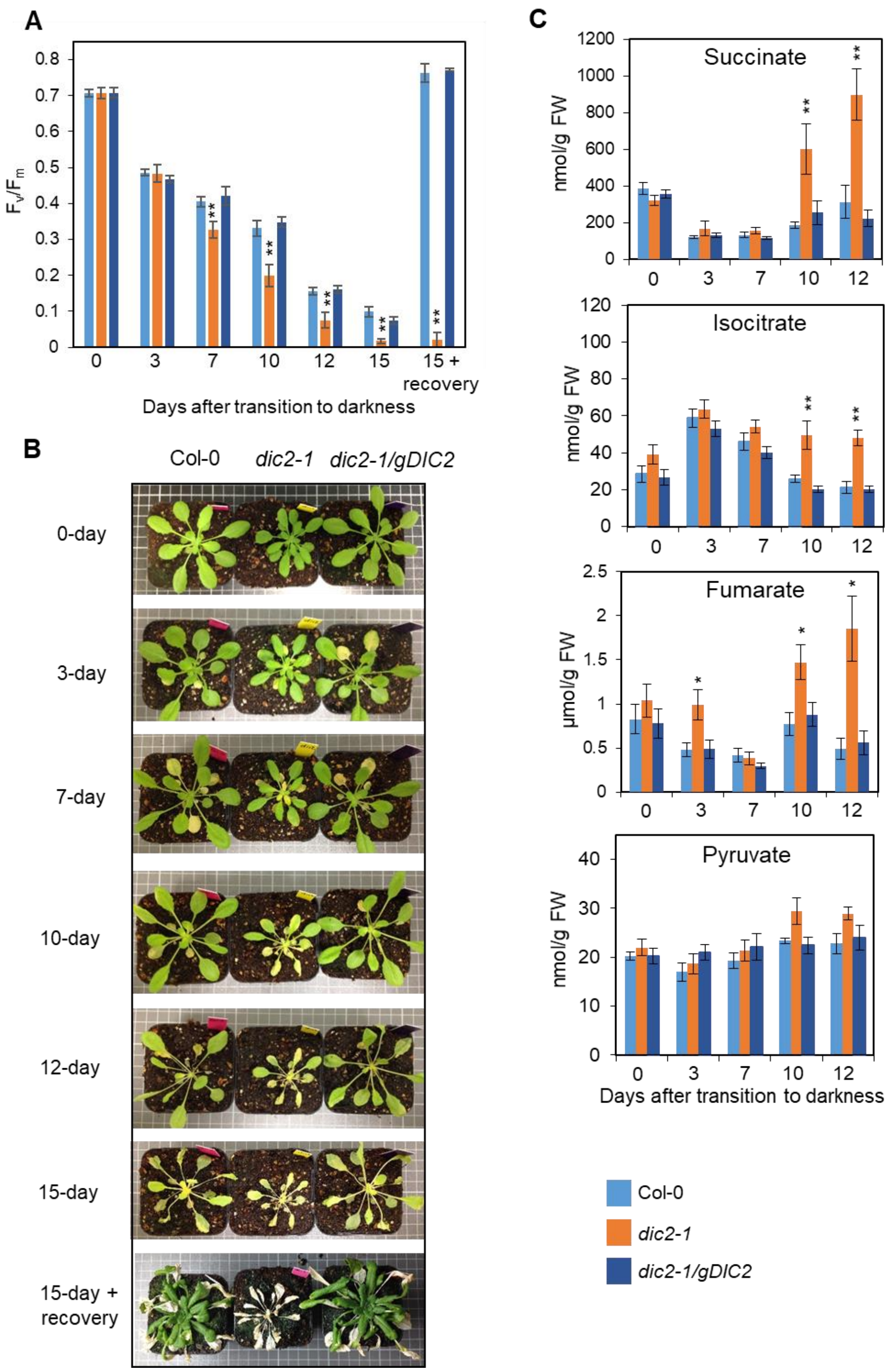


**Figure S8. Effects of extended dark treatment on Arabidopsis phenotypes.** (A) The maximum quantum efficiency of photosystem II ($F_v/F_m$) of five to six weeks old, short day grown plants treated with 0, 3, 7, 10, 12 and 15 days of extended darkness, as well as 15 days of extended darkness followed by 7 days of recovery phase in short day (n = 7). (B) Images of five to six weeks old, short-day grown plants treated with 0, 3, 7, 10, 12 and 15 days of extended darkness. (C) The abundances of succinate, isocitrate, fumarate and pyruvate in plants during extended dark treatment as measured by LC-SRM-MS. Values presented here are average ± S.E. (n = 7). Asterisks indicate a significant change as determined by the one-way ANOVA with Tukey post-hoc test (* $p < 0.05$; ** $p < 0.01$).

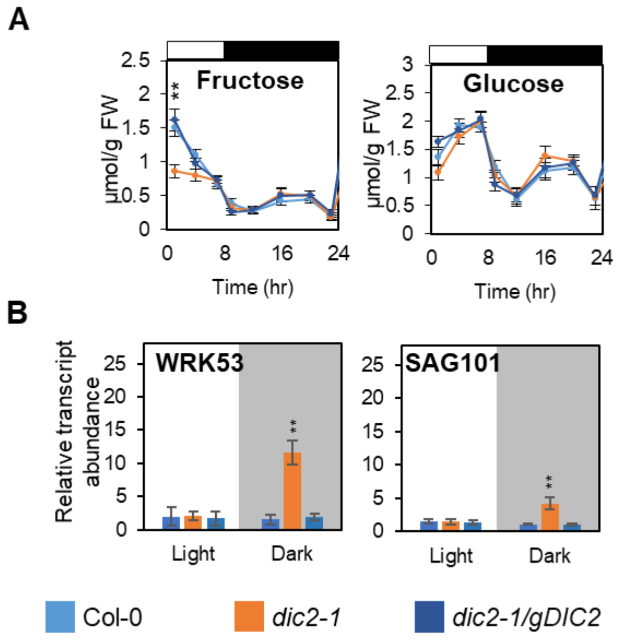


**Figure S9. The role of DIC2 in controlling sugar use in the light and dark.** (A) Fructose and glucose levels in leaf discs from Col-0, *dic2-1* and *dic2-1/gDIC2* collected at different time points of a diurnal cycle (see Figure 3 legend) as quantitatively determined by GC-MS against authentic standards (n = 8). (B) qPCR analysis showing the expression of SAG101 and WRKY53 in plants collected at the end of night (Shaded) or at the end of day (Light). All expression values were normalised against Col-0 end of day sample (n = 4). Data shown are mean ± S.E., and asterisks indicate a significant change as determined by the one-way ANOVA with Tukey post-hoc test (* $p < 0.05$; ** $p < 0.01$).

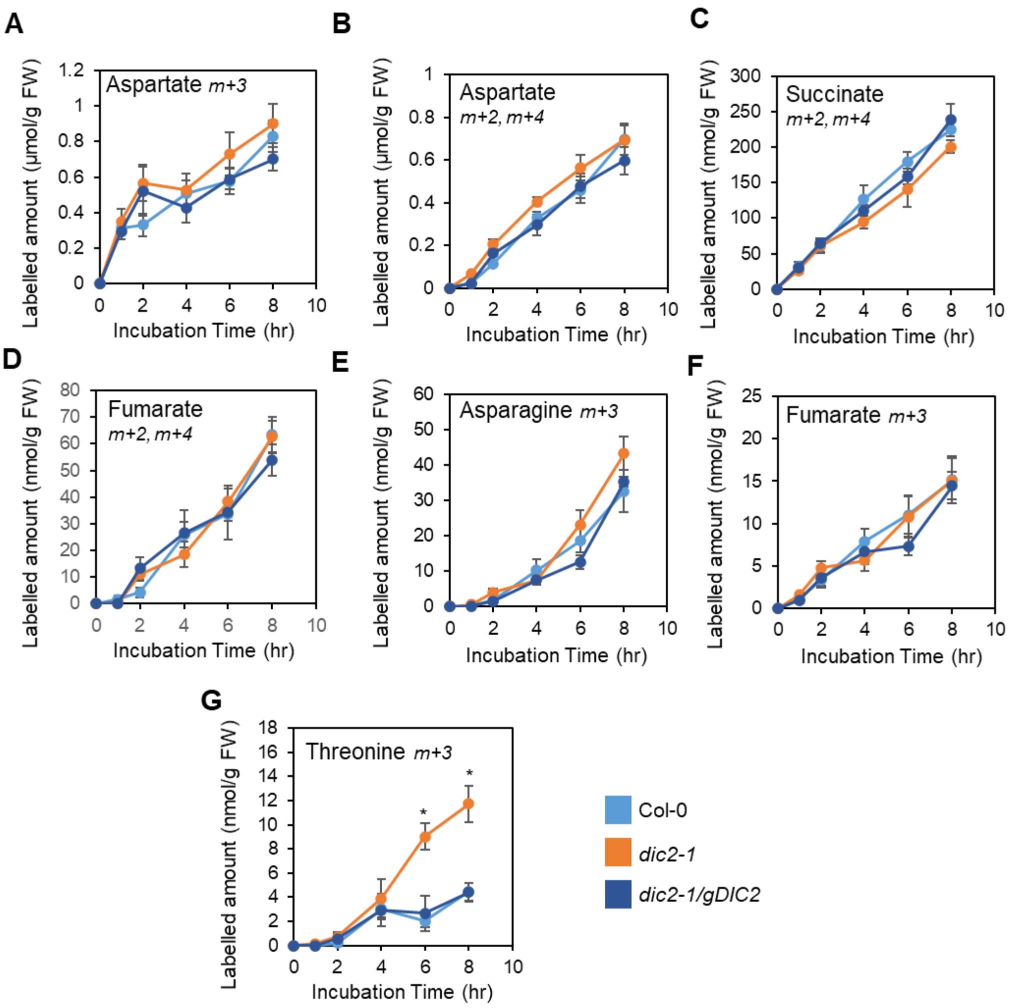


**Figure S10. Incorporation of $^{13}$C-glucose into metabolism of leaf discs in the dark.** Time-courses of $^{13}$C-labelling into metabolism in darkened leaf discs. Absolute abundance of labelled PEP-derived aspartate (A), TCA-derived aspartate (B), TCA-derived succinate (C), TCA-derived fumarate (D), PEP-derived asparagine (E), PEP-derived fumarate (F) and PEP-derived threonine (G). Means ± S.E. (n = 4). Asterisks indicating significant differences (* $p < 0.05$) as determined by one-way ANOVA Tukey post-hoc analysis.

```
DIC1      MGLKGFAEGGIASIVAGCSTHPLDLIKVRMQLQGESAPI--QTNLRPALAFQTSTTV---     55
DIC2      MGVKSFVEGGIASVIAGCSTHPLDLIKVRLQLHGEAPSTTTVTLLRPALAFPNSSPAAFL     60
          **:*.*.******::**************:**:**:      * ******* .*: .
                                        ←──── M ────→

DIC1      --NAPPLRVGVIGVGSRLIREEGMRALFSGVSATVLRQTLYSTTRMGLYDIIKGEWTDPE     113
DIC2      ETTSSVPKVGPISLGINIVKSEGAAALFSGVSATLLRQTLYSTTRMGLYEVLKNKWTDPE     120
            .:   :** *.:* .:::.**  *********:**************:::*.:*****
          ←── M ──→

DIC1      TKTMPLMKKIGAGAIAGAIGAAVGNPADVAMVRMQADGRLPLTDRRNYKSVLDAITQMIR     173
DIC2      SGKLNLSRKIGAGLVAGGIGAAVGNPADVAMVRMQADGRLPLAQRRNYAGVGDAIRSMVK     180
          : .: * :***** :**.************************::**** .* *** .*::
                                                ←──── M ────→

DIC1      GEGVTSLWRGSSLTINRAMLVTSSQLASYDSVKETILEKGLLKDGLGTHVSASFAAGFVA     233
DIC2      GEGVTSLWRGSALTINRAMIVTAAQLASYDQFKEGILENGVMNDGLGTHVVASFAAGFVA     240
          ***********:*******:**::******..** ***:*:::******* *********
          ← M →

DIC1      SVASNPVDVIKTRVMNMKVVAGVAPPYKGAVDCALKTVKAEGIMSLYKGFIPTVSRQAPF     293
DIC2      SVASNPVDVIKTRVMNMKVG-----AYDGAWDCAVKTVKAEGAMALYKGFVPTVCRQGPF     295
          *******************       *.** ***:******* *:*****:***.**.**
                          ←──── M ────→

DIC1      TVVLFVTLEQVKKLFKDYDF     313
DIC2      TVVLFVTLEQVRKLLRDF--     313
          ***********:**::*:
```

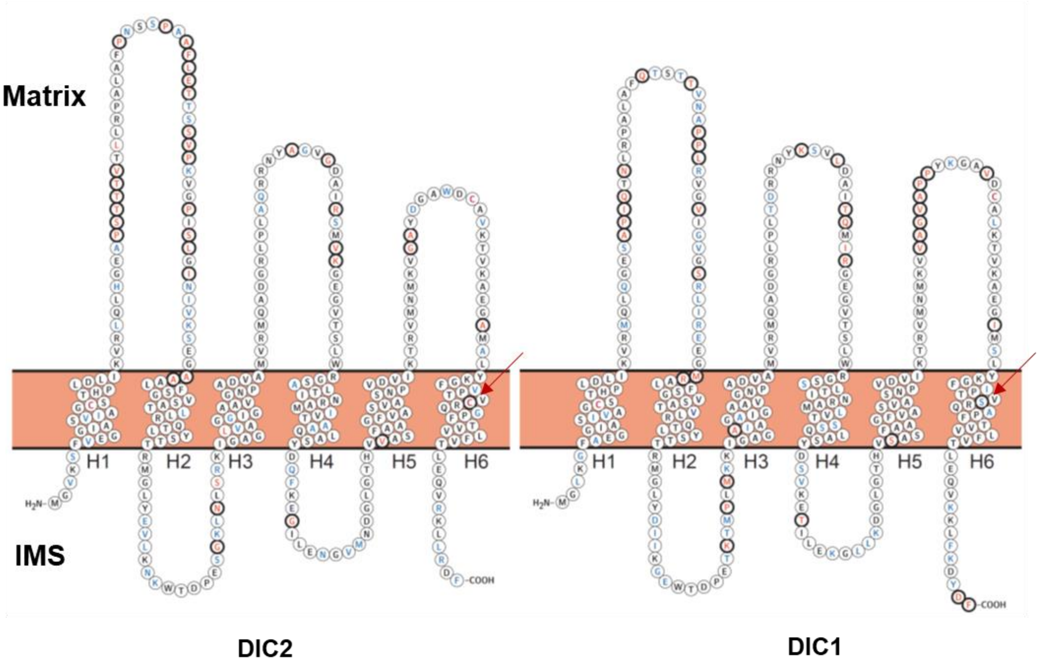


**Figure S11. Amino acid sequence comparison between DIC1 and DIC2.** *Upper panel*, sequence alignment of DIC1 and DIC2 was carried out by Clustal Omega. Cysteine residues are highlighted in red. Identical and similar amino acids are indicated by asterisk (*) and colon or dot (";" for conservation between groups of strongly similar properties and "." for conservation between groups of weakly similar properties), respectively. Predicted transmembrane domains are shown as blue boxes and predicted α-helices in the matrix are labelled as "M". *Lower panel*, cartoon representation of the predicted topology for DIC1 and DIC2. Each circle represents an individual amino acid. The location of cysteine residue unique to DIC2 is pointed by a red arrow. Differences in amino acid residues are highlighted in bold-black. The majority of differences in amino acid residues are found in the matrix side of both proteins.

#### Supplementary Tables

**Table S1. Relative expression of DIC2, DIC1 and DIC3 in Col-0, *dic2-1* and complemented lines as determined by qPCR.** Col-0 expression data were set to 1 and all other expression values were normalised against Col-0. n.d., expression not determined.

| | Relative expression | | |
|---|---|---|---|
| | **DIC2** | **DIC1** | **DIC3** |
| **Col-0** | 1 | 1 | 1 |
| ***dic2-1*** | 0 | 2 | 1 |
| ***dic2-1/gDIC2*** | 5 | 1 | 1 |
| ***dic2-1/DIC2 OE1*** | 440 | n.d. | n.d. |
| ***dic2-1/DIC2 OE2*** | 19 | n.d. | n.d. |
| ***dic2-1/DIC2 OE3*** | 1759 | n.d. | n.d. |
| ***dic2-1/DIC1 OE1*** | 0 | 3 | n.d. |
| ***dic2-1/DIC1 OE2*** | 0 | 5 | n.d. |
| ***dic2-1/DIC1 OE3*** | 0 | 10 | n.d. |
| ***dic2-1/DIC3 OE1*** | 0 | n.d. | 829 |
| ***dic2-1/DIC3 OE2*** | 0 | n.d. | 7 |
| ***dic2-1/DIC3 OE3*** | 0 | n.d. | 196 |
| ***DIC2 OE1*** | 500 | n.d. | n.d. |
| ***DIC2 OE2*** | 1531 | n.d. | n.d. |
| ***DIC2 OE3*** | 24 | n.d. | n.d. |
| ***DIC2 OE4*** | 45 | n.d. | n.d. |
| ***DIC2 OE5*** | 241 | n.d. | n.d. |

**Table S2. Kinetic properties of TCA cycle enzymes in isolated mitochondria.** Data expressed as mean ± S.E. from at least four replicates.

| | $K_m$ | | | | $V_{max}$ | | |
|---|---|---|---|---|---|---|---|
| | Col-0 | *dic2-1* | *dic2-1/gDIC2* | | Col-0 | *dic2-1* | *dic2-1/gDIC2* |
| **PDH (µM)** | 86 ± 11 | 90 ± 13 | 87 ± 15 | PDH (nmol/min/mg) | 114 ± 4 | 120 ± 5 | 126 ± 5 |
| **CS (µM)** | 28 ± 6 | 21 ± 4 | 28 ± 5 | CS (nmol/min/mg) | 292 ± 16 | 288 ± 14 | 284 ± 12 |
| **ACON (mM)** | 0.15 ± 0.02 | 0.14 ± 0.02 | 0.16 ± 0.09 | ACON (nmol/min/mg) | 194 ± 5 | 209 ± 5 | 201 ± 8 |
| **IDH (mM)** | 0.58 ± 0.08 | 0.67 ± 0.11 | 0.62 ± 0.09 | IDH (nmol/min/mg) | 147 ± 8 | 139 ± 5 | 135 ± 5 |
| **OGDH (µM)** | 33 ± 6 | 39 ± 6 | 34 ± 6 | OGDH (nmol/min/mg) | 140 ± 6 | 150 ± 8 | 133 ± 9 |
| **FUM (mM)** | 13 ± 4 | 10 ± 3 | 8.4 ± 2.7 | FUM (nmol/min/mg) | 261 ± 30 | 265 ± 25 | 215 ± 23 |
| **MDH (mM)** | 0.18 ± 0.04 | 0.15 ± 0.02 | 0.15 ± 0.03 | MDH (µmol/min/mg) | 14.2 ± 0.7 | 14.1 ± 0.7 | 13.8 ± 0.8 |
| **NAD-ME (mM)** | 0.42 ± 0.06 | 0.41 ± 0.03 | 0.38 ± 0.03 | ME (nmol/min/mg) | 294 ± 15 | 309 ± 7 | 282 ± 8 |

### Table S3. List of LC-MRM-MS transitions of peptides

| AGI number \| Protein Name | Peptide Sequence | Precursor m/z | Precursor Charge | Product Mz | Product Charge | Fragment Ion | Retention Time |
|---|---|---|---|---|---|---|---|
| AT3G01280\|VDAC1 | EDLIASLTVNDK | 659.348457 | 2 | 847.451973 | 1 | y8 | 8.57 |
| AT3G01280\|VDAC1 | EDLIASLTVNDK | 659.348457 | 2 | 776.414859 | 1 | y7 | 8.56 |
| AT3G01280\|VDAC1 | EDLIASLTVNDK | 659.348457 | 2 | 576.298767 | 1 | y5 | 8.56 |
| AT3G01280\|VDAC1 | FSITTFSPAGVAIT STGTK | 943.498923 | 2 | 1336.710707 | 1 | y14 | 9.9 |
| AT3G01280\|VDAC1 | FSITTFSPAGVAIT STGTK | 943.498923 | 2 | 1102.610265 | 1 | y12 | 9.9 |
| AT3G01280\|VDAC1 | FSITTFSPAGVAIT STGTK | 943.498923 | 2 | 1189.642293 | 1 | y13 | 9.9 |
| AT3G01280\|VDAC1 | INAGLSFTK | 475.768917 | 2 | 837.446494 | 1 | y8 | 6.6 |
| AT3G01280\|VDAC1 | INAGLSFTK | 475.768917 | 2 | 723.403566 | 1 | y7 | 6.6 |
| AT3G01280\|VDAC1 | INAGLSFTK | 475.768917 | 2 | 652.366452 | 1 | y6 | 6.6 |
| AT3G01280\|VDAC1 | SFFTISGEVDTK | 665.8299 | 2 | 848.435989 | 1 | y8 | 8.44 |
| AT3G01280\|VDAC1 | SFFTISGEVDTK | 665.8299 | 2 | 949.483667 | 1 | y9 | 8.44 |
| AT3G01280\|VDAC1 | SFFTISGEVDTK | 665.8299 | 2 | 735.351925 | 1 | y7 | 8.43 |
| At4g24570.1\|MCC | NYAGVGDAIR | 518.26453 | 2 | 461.243066 | 2 | y9 | 8.59 |
| At4g24570.1\|MCC | VGPISLGINIVK | 605.381905 | 2 | 527.336967 | 2 | y10 | 9.87 |
| At4g24570.1\|MCC | VGPISLGINIVK | 605.381905 | 2 | 555.847699 | 2 | y11 | 9.87 |
| At4g24570.1\|MCC | VGPISLGINIVK | 605.381905 | 2 | 643.413737 | 1 | y6 | 9.86 |
| AT4G26910.1\|PDH_DL ST | AAVSALQHQPVV NAVIDGDDIIYR | 855.45376 | 3 | 1079.536765 | 1 | y9 | 9.45 |
| AT4G26910.1\|PDH_DL ST | GLVVPVIR | 426.786913 | 2 | 484.324194 | 1 | y4 | 8.81 |
| AT4G26910.1\|PDH_DL ST | EAVYFLR | 449.245078 | 2 | 598.334758 | 1 | y4 | 7.87 |
| AT1G54220.1\|PDH | APALDYVDIPHSQ IR | 565.633533 | 3 | 737.405297 | 1 | y6 | 7.85 |
| AT3G13930.1\|PDH | IFASPLAR | 437.760895 | 2 | 614.362036 | 1 | y6 | 7.3 |
| AT3G13930.1\|PDH | VPALDYVDIPHTQ IR | 579.649183 | 3 | 751.420948 | 1 | y6 | 8.57 |
| AT3G13930.1\|PDH | GYIETPESMLL | 626.810121 | 2 | 689.353839 | 1 | y6 | 11.16 |
| AT3G52200.1\|PDH | NVPATIEGGR | 507.272355 | 2 | 800.426093 | 1 | y8 | 4.62 |
| AT3G52200.1\|PDH | MNVTLSADHR | 381.857357 | 3 | 312.177864 | 1 | y2 | 4.85 |
| AT3G52200.1\|PDH | IFDGQVGASFMS ELR | 828.906147 | 2 | 997.477142 | 1 | y9 | 10.12 |
| AT1G48030.1\|PDH_DL D | LGSEVTVVEFAG DIVPSMDGEIR | 807.405307 | 3 | 904.419293 | 1 | y8 | 11.75 |
| AT1G48030.1\|PDH_DL D | FPFMANSR | 485.234187 | 2 | 725.33992 | 1 | y6 | 7.83 |
| AT3G17240.1\|PDH_DL D | LGSEVTVVEFAA DIVPAMDGEIR | 806.745552 | 3 | 888.424378 | 1 | y8 | 13.14 |
| AT1G59900.1\|PDH_E1 | SVESSSQELLDFF R | 548.601898 | 3 | 584.282723 | 1 | y4 | 11.29 |
| AT1G59900.1\|PDH_E1 | GGSLHEVFSELM GR | 506.917164 | 3 | 363.1809 | 1 | y3 | 9.57 |
| AT1G59900.1\|PDH_E1 | GFGTESFGPDR | 585.264727 | 2 | 678.320565 | 1 | y6 | 6.3 |
| AT1G24180.1\|PDH_IA R1 | SVETSSEEILAFF R | 538.93851 | 3 | 653.376957 | 1 | y5 | 11.65 |
| AT1G24180.1\|PDH_IA R1 | NGPIILEMDTYR | 711.358301 | 2 | 927.424044 | 1 | y7 | 9.17 |
| AT5G50850.1\|PDH_M AB1 | SNYMSAGQINVPI VFR | 599.310173 | 3 | 631.392608 | 1 | y5 | 10.36 |
| AT5G50850.1\|PDH_M AB1 | LAEEGISAEVINLR | 757.414662 | 2 | 901.510157 | 1 | y8 | 8.86 |
| AT5G50850.1\|PDH_M AB1 | LALPQIEDIVR | 633.874445 | 2 | 485.271824 | 2 | y8 | 10.58 |
| AT2G44350.1\|CS4 | VPVVAAYVYR | 568.826766 | 2 | 742.38825 | 1 | y6 | 7.98 |
| AT2G44350.1\|CS4 | VIPGYGHGVLR | 389.892618 | 3 | 478.259051 | 2 | y9 | 5.95 |
| AT2G44350.1\|CS4 | LYEVVPPVLTELG K | 778.950349 | 2 | 953.566609 | 1 | y9 | 11.36 |
| AT2G05710.1\|ACO1 | VVNFSFDGQPAE LK | 775.896105 | 2 | 557.329339 | 1 | y5 | 8.63 |
| AT2G05710.1\|ACO1 | LSVFDAAMR | 505.260402 | 2 | 710.329021 | 1 | y6 | 8.38 |
| AT2G05710.1\|ACO1 | SSGEDTIILAGAE YGSGSSR | 978.96089 | 2 | 970.422464 | 1 | y10 | 7.7 |

| | | | | | | | |
|---|---|---|---|---|---|---|---|
| AT2G05710.1\|ACO1 | ILDWENTSTK | 603.803685 | 2 | 980.431966 | 1 | y8 | 6.85 |
| AT2G05710.1\|ACO1 | FSYNGQPAEIK | 627.311677 | 2 | 557.329339 | 1 | y5 | 5.78 |
| AT2G05710.1\|ACO1 | GVISEDFNSYGSR | 715.830962 | 2 | 1161.480707 | 1 | y10 | 6.61 |
| ICDH_AT3G09810.1 | ENTEGEYSGLEHQVVK | 606.955 | 3 | 673.340966 | 2 | y12 | 5.62 |
| ICDH_AT3G09810.1 | YPEIYYEK | 552.76604 | 2 | 471.234376 | 2 | y7 | 6.29 |
| ICDH_AT3G09810.1 | TADLGGSSTTTDFTK | 751.354467 | 2 | 1101.505859 | 1 | y11 | 5.5 |
| ICDH_AT5G03290.1 | VFTTAGVPIEWEEHYVGTEIDPR | 882.434254 | 3 | 985.468154 | 2 | y16 | 9.91 |
| ICDH_AT5G03290.1 | ENTEGEYSGLEHQVVR | 616.290383 | 3 | 687.34404 | 2 | y12 | 5.73 |
| ICDH_AT5G03290.1 | TDGLFLK | 397.226354 | 2 | 577.370809 | 1 | y5 | 7.44 |
| ICDH_AT5G14590.1 | LILPYLDLDIK | 658.397221 | 2 | 976.534974 | 1 | y8 | 12.34 |
| ICDH_AT5G14590.1 | VTVESAEAALK | 559.308603 | 2 | 818.425424 | 1 | y8 | 6.34 |
| ICDH_AT5G14590.1 | LIDDMVAYAVK | 619.328481 | 2 | 1011.481559 | 1 | y9 | 8.73 |
| ICDH1_AT4G35260.1 | VPPEVMESIR | 578.805173 | 2 | 529.270966 | 2 | y9 | 7.04 |
| ICDH1_AT4G35260.1 | TPVGGGVSSLNVQLR | 495.27909 | 3 | 629.372935 | 1 | y5 | 7.73 |
| ICDH1_AT4G35260.1 | DLGGTSTTQEVVDAVIAK | 601.982667 | 3 | 715.434866 | 1 | y7 | 9.44 |
| ICDH2_AT2G17130.1 | SLPEGLLESIK | 593.339903 | 2 | 493.281857 | 2 | y9 | 9.66 |
| ICDH2_AT2G17130.1 | TPVGGGVSSLNVNLR | 735.407172 | 2 | 1172.638211 | 1 | y12 | 7.68 |
| ICDH2_AT2G17130.1 | NANPVALLLSSAMMLR | 567.644387 | 3 | 795.385156 | 1 | y7 | 15.04 |
| AT3G55410.1\|OGDH_E1 | FGLEGGESLIPGMK | 717.868502 | 2 | 432.227516 | 1 | y4 | 10.05 |
| AT3G55410.1\|OGDH_E1 | AADLGVESIVIGMSHR | 552.290889 | 3 | 587.27184 | 1 | y5 | 9.61 |
| AT3G55410.1\|OGDH_E1 | YLQMSDDNPYVIPDMEPTMR | 805.693694 | 3 | 976.422663 | 1 | y8 | 9.18 |
| AT3G55410.1\|OGDH_E1 | LDPLGLEK | 442.758018 | 2 | 328.702514 | 2 | y6 | 7.53 |
| AT3G55410.1\|OGDH_E1 | FGLEGAESLIPGMK | 724.876327 | 2 | 432.227516 | 1 | y4 | 10.57 |
| AT3G55410.1\|OGDH_E1 | SADLGVENIVIGMPHR | 569.966406 | 3 | 597.292576 | 1 | y5 | 9.11 |
| AT2G20420.1\|succinyl_CoA_synthetase | AIQDVFPNESELVVK | 844.448702 | 2 | 1014.546602 | 1 | y9 | 9.28 |
| AT2G20420.1\|succinyl_CoA_synthetase | VPIDVFAGITDEDAAK | 554.285798 | 3 | 749.331189 | 1 | y7 | 10.09 |
| AT2G20420.1\|succinyl_CoA_synthetase | LITADDLDDAAEK | 695.340829 | 2 | 1163.506253 | 1 | y11 | 6.64 |
| AT2G20420.1\|succinyl_CoA_synthetase | GGTEHLGLPVFNSVAEAK | 609.319614 | 3 | 865.441408 | 1 | y8 | 13.32 |
| AT5G23250\|succinyl_CoA_synthetase | IGAAMFELFQER | 706.355562 | 2 | 968.483607 | 1 | y7 | 11 |
| AT5G66760.1\|SDH1_1 | SSYTIVDHTYDAVVGAGGAGLR | 770.057118 | 3 | 658.363098 | 1 | y8 | 8.73 |
| AT5G66760.1\|SDH1_1 | AVIELENYGLPFSR | 804.42503 | 2 | 506.272158 | 1 | y4 | 10.41 |
| AT5G66760.1\|SDH1_1 | SSQTILATGGYGR | 655.83859 | 2 | 794.415528 | 1 | y8 | 6.02 |
| AT5G66760.1\|SDH1_1 | SMTMEIR | 434.206781 | 2 | 649.333772 | 1 | y5 | 5.86 |
| AT5G66760.1\|SDH1_1 | TIAWLDR | 437.742702 | 2 | 660.346386 | 1 | y5 | 7.88 |
| AT5G66760.1\|SDH1_1 | IMQNNAAVFR | 582.303132 | 2 | 919.47444 | 1 | y8 | 6.05 |
| AT3G27380.1\|SDH2 | NEMDPSLTFR | 605.282062 | 2 | 720.403901 | 1 | y6 | 7.4 |
| AT1G47420.1\|SDH5 | VTVLGTSGLSGSYVEQR | 584.975735 | 3 | 531.288536 | 1 | y4 | 7.74 |
| At1G08480.1\|SDH6 | LSFFENYTR | 588.787838 | 2 | 829.383893 | 1 | y6 | 8.6 |
| At1G08480.1\|SDH6 | FMEWWER | 542.239469 | 2 | 805.362764 | 1 | y5 | 9.24 |
| At3G47833.1\|SDH7 | ALLAEDASLR | 529.795663 | 2 | 761.378808 | 1 | y7 | 6.91 |
| AT2G47510\|FUM1 | AAEILGR | 365.21632 | 2 | 587.351137 | 1 | y5 | 4.88 |
| AT2G47510\|FUM1 | AAEILGR | 365.21632 | 2 | 458.308544 | 1 | y4 | 4.88 |
| AT2G47510\|FUM1 | AAEILGR | 365.21632 | 2 | 345.22448 | 1 | y3 | 4.88 |
| AT2G47510\|FUM1 | AIMQAAQEVAEGK | 673.342651 | 2 | 404.213974 | 1 | y4 | 6.51 |
| AT2G47510\|FUM1 | AIMQAAQEVAEGK | 673.342651 | 2 | 902.457787 | 1 | y9 | 6.51 |
| AT2G47510\|FUM1 | AIMQAAQEVAEGK | 673.342651 | 2 | 831.420673 | 1 | y8 | 6.51 |

| | | | | | | | |
|---|---|---|---|---|---|---|---|
| AT2G47510\|FUM1 | EAALNLGVLTAEE FDTLVVPEK | 786.752812 | 3 | 373.208161 | 1 | y3 | 12.67 |
| AT2G47510\|FUM1 | EAALNLGVLTAEE FDTLVVPEK | 786.752812 | 3 | 472.276575 | 1 | y4 | 12.67 |
| AT2G47510\|FUM1 | EAALNLGVLTAEE FDTLVVPEK | 786.752812 | 3 | 571.344989 | 1 | y5 | 12.67 |
| AT2G47510\|FUM1 | SLQNFEIGGER | 625.312208 | 2 | 660.33113 | 1 | y6 | 6.92 |
| AT2G47510\|FUM1 | SLQNFEIGGER | 625.312208 | 2 | 418.204472 | 1 | y4 | 6.91 |
| AT2G47510\|FUM1 | SLQNFEIGGER | 625.312208 | 2 | 921.442471 | 1 | y8 | 6.92 |
| AT1G53240.1\|MDH1 | VAILGAAGGIGQP LALLMK | 897.039507 | 2 | 970.575399 | 1 | y9 | 12.79 |
| AT1G53240.1\|MDH1 | SEVVGYMGDDNL AK | 749.348131 | 2 | 1083.477536 | 1 | y10 | 6.69 |
| AT1G53240.1\|MDH1 | ALEGADLVIIPAG VPR | 795.964322 | 2 | 596.351471 | 1 | y6 | 10.5 |
| AT1G53240.1\|MDH1 | NGVEEVLDLGPL SDFEK | 930.964913 | 2 | 892.441074 | 1 | y8 | 11.89 |
| AT1G53240.1\|MDH1 | EGLEALKPELK | 409.571211 | 3 | 486.292225 | 1 | y4 | 6.93 |
| AT3G15020.1\|MDH2 | VVILGAAGGIGQP LSLLMK | 613.037502 | 3 | 801.490273 | 1 | y7 | 12.64 |
| AT3G15020.1\|MDH2 | SQVSGYMGDDDL GK | 736.322113 | 2 | 1157.47793 | 1 | y11 | 5.76 |
| AT3G15020.1\|MDH2 | NLSIAIAK | 415.260728 | 2 | 602.387188 | 1 | y6 | 7.07 |
| At2g13560.1\|NAD-ME1 | IAAAVIK | 343.233982 | 2 | 572.376623 | 1 | y6 | 5.11 |
| At2g13560.1\|NAD-ME1 | IAAAVIK | 343.233982 | 2 | 501.339509 | 1 | y5 | 5.11 |
| At2g13560.1\|NAD-ME1 | IAAAVIK | 343.233982 | 2 | 430.302396 | 1 | y4 | 5.11 |
| At2g13560.1\|NAD-ME1 | IVVAGAGSAGIGV LNAAR | 532.645772 | 3 | 431.236107 | 1 | y4 | 8.43 |
| At2g13560.1\|NAD-ME1 | IVVAGAGSAGIGV LNAAR | 532.645772 | 3 | 544.320171 | 1 | y5 | 8.43 |
| At2g13560.1\|NAD-ME1 | IVVAGAGSAGIGV LNAAR | 532.645772 | 3 | 700.410049 | 1 | y7 | 8.42 |
| At2g13560.1\|NAD-ME1 | WPHVIVQFEDFQ SK | 587.298261 | 3 | 753.34136 | 1 | y6 | 9.41 |
| At2g13560.1\|NAD-ME1 | WPHVIVQFEDFQ SK | 587.298261 | 3 | 624.298767 | 1 | y5 | 9.4 |
| At2g13560.1\|NAD-ME1 | WPHVIVQFEDFQ SK | 587.298261 | 3 | 900.409774 | 1 | y7 | 9.4 |
| At4g00570.1\|NAD-ME2 | AVVQFEDFQAK | 641.327327 | 2 | 884.414859 | 1 | y7 | 7.57 |
| At4g00570.1\|NAD-ME2 | AVVQFEDFQAK | 641.327327 | 2 | 737.346445 | 1 | y6 | 7.56 |
| At4g00570.1\|NAD-ME2 | AVVQFEDFQAK | 641.327327 | 2 | 1012.473437 | 1 | y8 | 7.57 |
| At4g00570.1\|NAD-ME2 | GILYPSINNIR | 630.358962 | 2 | 976.521056 | 1 | y8 | 8.44 |
| At4g00570.1\|NAD-ME2 | GILYPSINNIR | 630.358962 | 2 | 813.457727 | 1 | y7 | 8.43 |
| At4g00570.1\|NAD-ME2 | GILYPSINNIR | 630.358962 | 2 | 407.232502 | 2 | y7 | 8.44 |
| At4g00570.1\|NAD-ME2 | IVVVGAGSAGLG VTK | 664.400827 | 2 | 917.505071 | 1 | y11 | 7.42 |
| At4g00570.1\|NAD-ME2 | IVVVGAGSAGLG VTK | 664.400827 | 2 | 1016.573485 | 1 | y12 | 7.43 |
| At4g00570.1\|NAD-ME2 | IVVVGAGSAGLG VTK | 664.400827 | 2 | 789.446494 | 1 | y9 | 7.41 |

**Table S4. List of primers.**

| Name | Primer sequence | Purpose |
|---|---|---|
| DIC1_GW_Fwd | GGGGACAAGTTTGTACAAAAAAGCAGGCTAAAAAATGGGTCTAAAGGGTTTTGC | Generation of DIC1 construct by Gateway cloning |
| DIC1_GW2_Rev | GGGGACCACTTTGTACAAGAAAGCTGGGTCTCAAAAGTCATAGTCCTTGAACAAC | Generation of DIC1 construct by Gateway cloning |
| DIC2_GW_Fwd | GGGGACAAGTTTGTACAAAAAAGCAGGCTAAAAAATGGGAGTCAAAAGTTTCGTTG | Generation of DIC2 and DIC2-GFP construct by Gateway cloning |
| DIC2_GW_Rev | GGGGACCACTTTGTACAAGAAAGCTGGGTCAAAATCTCGAAGCAGCTTCCT | Generation of DIC2 construct by Gateway cloning |
| DIC2_GW2_Rev | GGGGACCACTTTGTACAAGAAAGCTGGGTCTCAAAAATCTCGAAGCAGCTTCCT | Generation of DIC2-GFP construct by Gateway cloning (stop deleted) |
| DIC3_GW_Fwd | GGGGACAAGTTTGTACAAAAAAGCAGGCTAAAAAATGGGCTTCAAACCATTTCTTG | Generation of DIC3 construct by Gateway cloning |
| DIC3_GW2_Rev | GGGGACCACTTTGTACAAGAAAGCTGGGTCTCAAAATTTAACGTCTTTTAACAGACC | Generation of DIC3 construct by Gateway cloning |
| DIC2_COMP2_Fwd | CCGGAATTCGAGCACAAGTTGGGTTTTCTATTGTTCTC | Generation of DIC2 construct for complementation, EcoRI cut |
| DIC2_COMP4_Rev | GTAAGGATCCGGACCTTCTCCAGATGATACTCTTT | Generation of DIC2 construct for complementation, BamHI cut |
| At4g24570_FWD | ATGGGAGTCAAAAGTTTCGTTGAAGGTGG | *mcc-1* and *mcc-2* genotyping |
| At4g24570_REV | CTAACTTGCTCCAACGTAACGAAGAGAAC | *mcc-1* and *mcc-3* genotyping |
| GABI_T-DNA | ATATTGACCATCATACTCATTGC | *mcc-1* genotyping |
| GK-833F11_LB | TTGTCTAACTCATAGCAACAAGTCG | *mcc-2* genotyping |
| CS66519_LP | GAGGTTTTCCCTGACAAGGAG | *mccb1-1* genotyping |
| CS66519_RP | AGACCCGAGCTTAGAATCTGG | *mccb1-1* genotyping |
| CS66518_LP | GAGCTTACTTTGCTTCGAAACC | *mcca1-1* genotyping |
| CS66518_RP | CAAAACTGAAACAAGGCCTTG | *mcca1-1* genotyping |
| CS66521_LP | GCACAGAGGAACGAAAACTTG | *hml1-2* genotyping |
| CS66521_RP | GAAGTTGGTCCAAGAGATGGC | *hml1-2* genotyping |
| gdh1-2_LP | GATCGCAAGAGTGAGTTTTGC | *gdh1-2* genotyping |
| gdh1-2_RP | TATGAACCAGGATCTTGGTGG | *gdh1-2* genotyping |
| gdh2-1_LP | TAGCAAGCCACACAACTGATG | *gdh2-1* genotyping |
| gdh2-1_RP | TGTAGTATTTGCCACCGAAGC | *gdh2-1* genotyping |
| SALK_LB | ATTTTGCCGATTTCGGAAC | *mccb1-1, mcca1-1, hml1-2, gdh dKO* genotyping |
| EF-1α-F | TGAGCACGCTCTTCTTGCTTTCA | qRT-PCR reference gene EF-1α |
| EF-1α-R | GGTGGTGGCATCCATCTTGTTACA | qRT-PCR reference gene EF-1α |
| SAND_F | CCATATTGCAAGAAGTTTGCGCGTCTG | qRT-PCR reference gene *Sand Family Protein* |
| SAND_R | GCAAGTCATCGGGATGGAGAGACG | qRT-PCR reference gene *Sand Family Protein* |
| DIC1_qPCR_F | AACAATTCAAAATGGGTCTAAAGG | qRT-PCR of DIC1 |
| DIC1_qPCR_R | CCGCAACAATCGAAGCTATT | qRT-PCR of DIC1 |
| DIC2C_qPCR_F | TGCGGTGAAGACGGTTAAA | qRT-PCR of DIC2 |
| DIC2C_qPCR_F | AAAGGACCTTGCCTACAAACTG | qRT-PCR of DIC2 |
| DIC3_qPCR_F | AACGAAGGTGGACTGATCAAC | qRT-PCR of DIC3 |
| DIC3_qPCR_R | AGGATTCCCGACGACTGAT | qRT-PCR of DIC3 |
| UCP1_qPCR_F | GGACTGGGAGCAGGATTCTT | qRT-PCR of UCP1 |
| UCP1_qPCR_R | TCCCATCATTCTTGACTTAACCA | qRT-PCR of UCP1 |

| | | |
|---|---|---|
| DTC_qPCR_F | CCGATCGACATGATTAAGGTG | qRT-PCR of DTC |
| DTC_qPCR_R | CACCAACGCCCTCATTCT | qRT-PCR of DTC |
| SFC_qPCR_F | CCATTGTTACACCCTTTGAGG | qRT-PCR of SFC |
| SFC_qPCR_R | CTTGAAAAGCTCAGGACTCAATC | qRT-PCR of SFC |
| SEN1_qPCR_F | CCTCAACTGATCTTCTCACTGC | qRT-PCR of SEN1 |
| SEN1_qPCR_R | TTCTCTGTCCAAGCGACGTA | qRT-PCR of SEN1 |
| Sag101_qPCR_F | TGTCTTCTCCACAGATCTATTCCAG | qRT-PCR of SAG101 |
| Sag101_qPCR_R | CTCCATGAGCTATGTAAGAGACCAG | qRT-PCR of SAG101 |
| WRKY53_F | CAGACGGGGATGCTACGG | qRT-PCR of WRKY53 |
| WRKY53_R | GGCGAGGCTAATGGTGGTG | qRT-PCR of WRKY53 |

#### Table S5. List of LC-MRM-MS transitions of metabolites

| Name | Precursor Ion | Product Ion 1 | Product Ion 2 | Product Ion 3 | Retention time (min) | Identification method | ESI mode |
|---|---|---|---|---|---|---|---|
| GABA | 104.1 | 87.1 | 45.2 | 69.2 | 0.84 | Authentic standard | + |
| Alanine | 90 | 44.1 | | | 0.95 | Authentic standard | + |
| Serine | 106.1 | 60.2 | 42.2 | | 0.96 | Authentic standard | + |
| Asparagine | 133.1 | 46.1 | 74.1 | | 0.96 | Authentic standard | + |
| Glutamine | 147.2 | 130 | 41.2 | 84.2 | 0.98 | Authentic standard | + |
| Threonine | 120.1 | 74 | 55.8 | | 0.98 | Authentic standard | + |
| Glutamate | 148.1 | 102.1 | 84.1 | | 1.03 | Authentic standard | + |
| Proline | 116.1 | 70 | 43.1 | | 1.04 | Authentic standard | + |
| Valine | 118.1 | 72.2 | 42.1 | 57.4 | 1.1 | Authentic standard | + |
| Aspartate | 134.1 | 88.1 | 74.1 | | 1.24 | Authentic standard | + |
| Methionine | 150.2 | 61.2 | 104.3 | | 1.29 | Authentic standard | + |
| Leucine/Isoleucine | 132.1 | 86.2 | 69.1 | | 1.45 | Authentic standard | + |
| 13C-Leucine (Internal standard) | 138.1 | 46.2 | | | 1.46 | Authentic standard | + |
| Tyrosine | 182.2 | 136.2 | 91.2 | 119.3 | 1.82 | Authentic standard | + |
| Phenylalanine | 166.2 | 120.1 | 77.2 | 118.1 | 2.51 | Authentic standard | + |
| Tryptophan | 205.2 | 146.2 | 118.1 | | 5.74 | Authentic standard | + |
| Malate | 403.3 | 208 | 137 | | 7.5 | Authentic standard | - |
| Succinate | 387.3 | 234 | 98 | 152.1 | 7.7 | Authentic standard | - |
| D-2-hydroxyglutarate | 417.1 | 137 | 264 | | 7.9 | Authentic standard | - |
| Adipic acid (Internal standard) | 415.2 | 178 | 150 | | 8.4 | Authentic standard | - |
| Fumarate | 385.3 | 232.1 | 96 | 152.1 | 8.5 | Authentic standard | - |
| Isocitrate | 596.4 | 234.1 | 387.1 | | 9.2 | Authentic standard | - |
| Citrate | 596.4 | 222.1 | 401.2 | | 9.5 | Authentic standard | - |
| Pyruvate | 357.3 | 137 | 150 | | 10.1 | Authentic standard | - |
| 2-oxoglutarate | 550.4 | 371.2 | 233.2 | | 10.7 | Authentic standard | - |
